# Joint decomposition of Hi-C maps reveals salient features of genome architecture across tissues and development

**DOI:** 10.1101/2025.08.07.669116

**Authors:** Thomas Reimonn, Vedat Yilmaz, Hoang Tran, Garrett Ng, Derek Liu, Nezar Abdennur

## Abstract

The spatial organization of chromosomes in the nucleus is fundamental to cellular processes. Contact frequency maps from Hi-C and related chromosome conformation capture assays are increasingly available for a wide variety of biosamples and conditions, creating opportunities for comprehensive studies of genome compartmentalization and long-range interactions. However, the conventional dimensionality reduction approach to study long-range contact frequency profiles projects individual datasets into different and incomparable linear subspaces, making the resulting embeddings unsuitable for large-scale integrative analysis. To address this shortcoming and overcome the computational constraints involved in doing so, we introduce an analytic framework and Python toolkit that leverages incremental principal component analysis to project interchromosomal contact frequency profiles across arbitrarily many Hi-C datasets onto a common set of components or basis vectors. Our approach produces robust and directly comparable first and higher-order principal component (PC) scores that collectively capture biologically meaningful information beyond traditional A/B compartments. By applying our framework to a collection of 89 human Hi-C samples, we uncover distinct patterns of nuclear architecture reflecting cell state categories, associated with different heterochromatin state compositions. We also demonstrate that jointly-derived higher-order PCs improve the prediction of gene expression and regulatory element activity during differentiation. Together, our joint decomposition approach provides a powerful and scalable foundation for systematically investigating genome organization, providing critical insights into its role in development and disease.

## Introduction

Chromosome organization plays a pivotal role in cellular processes, including gene regulation, organ development, and cell identity determination [1–3]. The principles that shape genome architecture are crucial for facilitating physical proximity between regulatory elements and target genes, as well as replicating and maintaining chromosomes, and transferring them during cell division [4,5]. The evolution of chromatin conformation capture-based molecular assays (e.g. Hi-C, Micro-C) has enabled comprehensive profiling of contact frequency maps of many biospecimens and conditions at multiple resolution scales [6–8]. Over the last two decades, these methods have revealed some of the major biophysical processes shaping genome architecture [9]. The first of these, loop extrusion by ATP-dependent SMC complexes, influences local genome organization in *cis* during interphase, giving rise to the patterns in contact maps collectively associated with Topologically Associating Domains (TADs) in vertebrates [10,11]. The second major process, termed *compartmentalization*, is mechanistically independent of loop extrusion and gives rise to a genome-wide checkering pattern between chromosome domains within the same and across different chromosomes [12–14].

The checkering pattern seen in most interphase Hi-C maps is usually characterized as a binary partition of two groups of loci that interact preferentially with each other. In mammalian cells, these two groups or “compartments” are referred to as A and B and broadly correspond to transcriptionally active and inactive loci, respectively. The conventional analysis of genome compartmentalization consists of the calculation of the leading eigenvector of a suitably pre-processed normalized contact matrix or its correlation matrix, obtained from either intra- or inter-chromosomal maps [13,15,16]. The mathematical reason this one-dimensional profile is taken to define A/B compartments is because its outer product produces the checkerboard-like matrix of rank 1 that best reconstructs the input matrix in a least-squares sense [17–19].

Studies using higher-resolution contact maps have long indicated that the A/B delineation and the continuous rank-1 quantification provide only an approximation of the complex patterns of long-range interactions [20,21]. The partitioning of genomic loci into greater than two groups based on long-range contact frequency profiles is often referred to as identifying “sub-compartments”, though we have proposed Interaction Profile Groups (IPGs) as a more neutral, less suggestive term. The first methodologies for IPG identification applied unsupervised clustering directly on interchromosomal contact matrices [20,22] and other heuristic approaches have been developed for A/B compartment and IPG identification from intrachromosomal contact matrices [23,24]. Polymer simulations based on mechanistic biophysical principles as well as constraint-based and generative modeling approaches based on mechanistically driven assumptions have subsequently provided insight into the drivers of compartmentalization [25–28]. Unsupervised dimensionality reduction provides an essential complement to mechanistic modeling, by enabling the appraisal of the full scope of long-range interactions and their variability across cell types and conditions, as well as distinguishing biological from technical source of variation. In fact, the traditional matrix decomposition approach to quantifying A/B compartments extends naturally to approximations of higher matrix rank (**Figure 1A**). By generating additional component vectors, the reconstruction of the input matrix is improved, thereby better explaining the variance in the data. We recently showed how unsupervised clustering on such higher-order latent representations provides a scalable method for elucidating IPGs in individual Hi-C samples [21].

**Figure 1.**
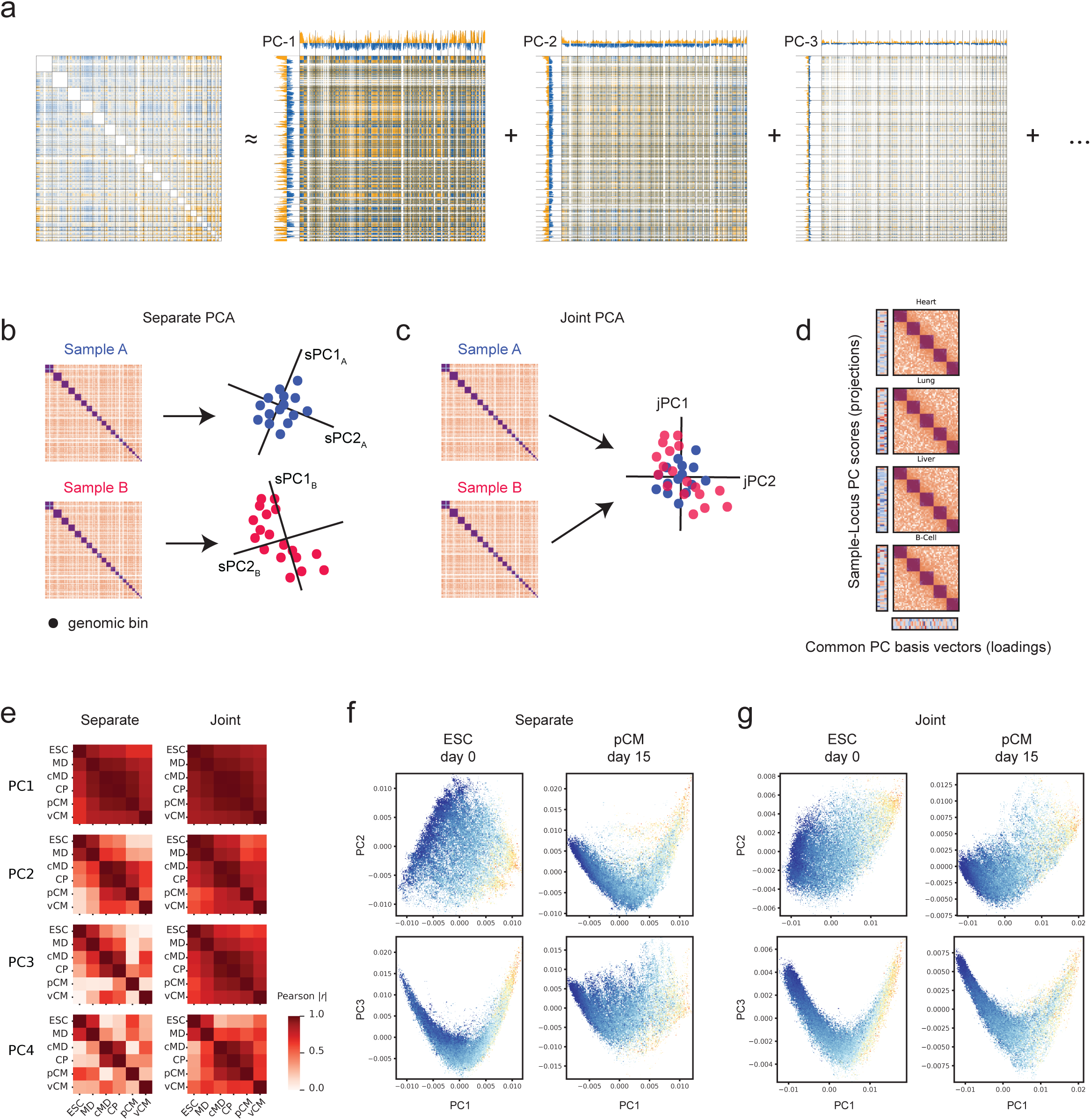
Joint PCA provides coherent, comparable embeddings of multiple Hi-C datasets. (A) Schematic of the factorization of a genome-wide contact frequency map (or a derived similarity matrix) by a truncated eigendecomposition after mean centering. The sum of the eigenvalue-weighted outer products of the first *n* eigenvectors (or principal components) gives the best rank-*n* reconstruction of the input matrix in terms of overall squared error. Traditional A/B compartment analysis corresponds to calculating only the first such component vector for a single contact matrix. (B) Cartoon of applying PCA to two Hi-C datasets separately, yielding incompatible principal component projections in distinct vector spaces. (C) Depiction of jointly applying PCA to produce a common set of *n* components (i.e. basis vectors) spanning a common vector space. Each sample dataset can be projected onto this coordinate system to produce *n* PC score vectors. Each sample-locus pair is thus associated with an *n*-dimensional compressed representation of its overall contact frequency profile. (D) Representation of joint PCA of multiple Hi-C datasets as performed by *jointly-hic* that produces sample-specific projections or embeddings (PC score vectors) onto a common set of basis vectors across all input samples. (E) Absolute Pearson correlation coefficients between score vectors derived by separate PCA (left) and by joint PCA (right) for six stages of an *in vitro* cardiomyocyte differentiation. (F-G) PC score scatter plots for PC1 vs PC2 (top) and PC1 vs PC3 (bottom) at the embryonic stem cell (ESC, left) and primitive cardiomyocyte (pCM, right) stages obtained by separate (F) and joint (G) PCA decompositions shows “switching” of sPC2 and sPC3 between stages but consistency between jPC2 and jPC3.

Overall, the origins and details of compartmentalization and other global features of genome organization in the nucleus still remain unclear. While specific biochemical affinities between chromatin states are thought to be a major driving mechanism of chromosome compartmentalization [9,29], evidence is also growing for the importance of the association of certain regions of the genome with nuclear bodies (e..g, speckles, nucleolus) and tethering to the nuclear lamina for understanding spatial and functional genome organization [30]. Indeed, classic A/B compartment intervals tend to be distributed radially in the nucleus, with euchromatin typically located centrally and inactive or heterochromatin located towards the nuclear periphery. Additionally, because of the slow timescale of global compartmental organization in interphase, Hi-C maps can also be influenced by the configurations of chromosomes upon exit from mitosis [31–33].

Advancements in the resolution and production of contact frequency maps, including from large concerted efforts such as ENCODE and the 4D Nucleome Consortium, has led to a diversity of 3D genomic datasets across a range of human cell types and tissues [34–37]. These datasets provide opportunities for integrative analyses across diverse biological contexts. However, current analytical frameworks are not yet suitable for the integration of many heterogeneous samples. The standard approach is to perform matrix decomposition separately on each sample, which results in genomic loci being embedded into different linear subspaces (**Figure 1B**). As a result, while the first component vector derived from each of two contact matrices might explain reasonably well the primary compartmentalization pattern in each sample, strictly speaking, those two vectors are not compatible for one-to-one comparison. Careful normalization and manual post-processing steps are required to guarantee sufficient compatibility of A/B scores to support differential analysis [38,39]. Overall this approach greatly limits what can be learned by integrating information from numerous contact frequency maps simultaneously.

Producing a *joint* embedding from all input samples avoids the incompatibility introduced by separate decomposition and avoids introducing biases. However, the computational limitations for memory and compute often make such large-scale joint analyses infeasible. To directly address this methodological gap, we present an integrative analytic framework and Python toolkit, called *jointly-hic*, that computes joint decompositions of interchromosomal contact frequency profiles from multiple datasets simultaneously, situating embeddings from all chromosomes and samples in a unified vector space (**Figure 1C, D**). The toolkit applies a mini-batch incremental principal component analysis (PCA) algorithm that scales to arbitrary input sizes without incurring additional memory costs [40]. We establish the effectiveness of our method to coherently project multiple Hi-C maps into a common space spanned by multiple biologically informative basis vectors. We applied *jointly-hic* to a diverse collection of 89 Hi-C datasets, encompassing primary cells, human *ex vivo* tissues, and *in vitro* differentiation models of heart, pancreas, and liver development [41,42]. Leveraging this atlas, we identified distinct nuclear organization patterns distinguishing immune cells, *in vitro-*derived cells, and other mature tissue samples, characterized in part by differences in repressive histone modifications. Finally, by integrating joint embeddings with gene expression and chromatin accessibility profiles, we show that changes in higher-order principal component scores are predictive of differential gene expression and regulatory element activity beyond the traditional A/B compartment score. Collectively, our approach establishes a scalable and robust foundation for systematically investigating genome organization across diverse biological contexts in order to provide critical insights into the structural underpinnings of gene regulation and cellular identity.

## Results

### Joint PCA produces robust and coherent embeddings of long-range genomic contact frequencies from multiple samples

We developed a framework that performs a joint PCA of interchromosomal contact frequency profiles from multiple Hi-C datasets simultaneously (**Figure 1C, D**). We begin by defining some terminology. The conventional matrix-based approaches to the dimensionality reduction of Hi-C data are mathematically equivalent to variants of PCA, treating the rows of the input matrix as observations or training examples (**see Methods**). From the matrix factorization perspective, one obtains a single collection of eigenvectors, whose weighted outer products sum to an approximation of the original matrix (**Figure 1A**). From the PCA perspective, one obtains a set of principal components (PCs) or *basis vectors* over the space of input features that define a new orthogonal coordinate system and a set of PC *score vectors* corresponding to the projections of the input observations onto the basis vectors. For a single symmetric contact matrix as input, the feature vectors (columns) and observations (rows) are the same and consequently the basis and score vectors are also identical. As such, the distinction between the two concepts has not been relevant to the field in practice. However, for a joint PCA trained on more than one input contact matrix, this distinction must be made explicitly. The principal component basis vectors are of cardinality the number of genomic bins or loci (features) and define a common set of coordinate axes in which to project the observations (**Figure 1C**). Each observation, a row of an input sample contact matrix, corresponds to the interaction profile of what we term a *sample-locus*, and is associated with projections or scores along each basis vector. Therefore, each input sample matrix is associated with its own set of PC score vectors, one for each corresponding basis vector (**Figure 1D**). Since we seek a minority of basis vectors that explain most of the variance, the linear projections of sample-locus interaction profiles map to a lower-dimensional feature space, so we also refer to them as embeddings.

We first sought to evaluate the robustness of our joint PCA approach to variation in data quality. To do this, we started with a deeply sequenced Hi-C dataset from the HCT116 human cell line and introduced synthetic perturbations [43]. We progressively downsampled the dataset up to 90% to simulate varying sequencing depths. We also added progressively increasing levels of random ligation noise to simulate poor library quality. Despite the data perturbations, comparisons of scatter plots of PC1 vs PC2 scores and the correlation map between PC score vectors indicate an excellent preservation of embedding similarity in the face of lower read depth and remarkable resilience despite substantial simulated increases in random ligation noise (**S.Figure 1A, 1B**). These results confirm that joint decomposition integrates Hi-C data across a broad depth and quality spectrum in a robust and consistent manner.

We next sought to explore the mutual compatibility of jointly-derived (jPCA) versus separately-derived (sPCA) principal component embeddings from multiple samples for comparative analysis across biological conditions. For this purpose, we considered a sequence of six developmental stages from a published *in vitro* cardiomyocyte differentiation study: embryonic stem cells (ESC), mesoderm (MD), cardiac mesoderm (cMD), cardiac progenitor (CP), primitive cardiomyocyte (pCM), and ventricular cardiomyocyte stages (vCM) [42]. For each of the 50-kb pre-processed contact matrices corresponding to a differentiation stage, we generated PC score vectors by both (i) sPCA of each stage’s contact matrix separately and (ii) jPCA of the stage’s contact matrix with those from all the other stages. We assessed the relationships between separately-derived score vectors (sPC) and jointly-derived score vectors (jPC) by calculating matrices of absolute Pearson correlation between score vectors and by inspecting scatter plots of genomic loci by PC scores (**Figure 1E-G, S.Figure 1C-F**).

We note that a jPCA embedding of multiple contact maps will produce a set of orthonormal basis vectors along which the variance explained by the profiles of genomic loci in all contact maps in the collection is maximized (**Figure 1C**). Hence, the exact directions of the common jPC basis vectors are a consensus or “compromise” that will depend on the nature of the variation within the samples that are used to construct the joint embedding [44]. Nevertheless, when combining interphase datasets, the first component is expected not to deviate from the general direction that explains A/B compartmentalization, as this is the dominant source of pattern variation in virtually all mammalian interphase maps and, despite cell type-specific variation, is generally robust and universal. Despite both sPC and jPC score vectors being strongly correlated across stages, jPC1 scores showed consistently stronger inter-stage correlation. Consistent with these observations, we observe that sPCA and jPCA both display a gradient of GC-content along the respective PC1 axis in all stages (**Figure 1F-G, S.Figure 1C, 1D**).

In contrast, the nature of the interaction patterns captured by higher-order jPCs is expected to be more strongly influenced by the specific composition of contact maps in the sample set. Indeed, we observed an immediate deterioration in inter-stage (absolute) Pearson correlation between higher-order sPC vectors, while inter-stage similarity remained high between jPC score vectors (**Figure 1E**). Comparing higher-order sPC to jPC vectors also indicated that sPC vectors of a given rank sometimes capture different sources of variation in different stages and may exhibit a high degree of correlation with sPC vectors of a different rank in another stage. As an illustrative example, in the first differentiation stage (ESC), the sPC1-sPC2 point cloud forms a characteristic “sail-shaped” pattern, while the sPC1-sPC3 appears “crescent-shaped”. Interestingly, by the third stage (cMD) onwards this relationship is inverted: sPC1-sPC2 takes on the crescent shape while sPC1-sPC3 looks like a sail. This inversion is reflected in the angles between the sPC2 and sPC3 basis vectors in different stages being as low as ∼47.6° (**S.Figure 1G**), implying that across stages basis vectors of different rank are linearly dependent rather than orthogonal as they are within the same stage.

Because the common set of jPC basis vectors are mutually orthogonal across all stages by construction, the qualitative appearance of the point clouds in the jPC score scatter plots remains consistent across all stages, with all jPC1-jPC2 and jPC1-jPC3 point clouds adopting sail and crescent appearances, respectively. Furthermore, gradients of GC content and distance from the centromere remain stable in the jPC scatter plots but often vary from stage to stage in the sPC scatters. Overall, our results suggest that while similar linear subspaces may explain the variance of the six contact maps in each sPCA embedding and the jPCA embedding, in sPCA the relative importance of the basis vectors can change from sample to sample, making, for example, sPC2 from embryonic stem cells more closely aligned with sPC3 than with than with sPC2 from cardiomyocytes and vice versa. On the other hand, jPCA ensures that higher-order PC scores are always comparable between samples.

Our results further revealed that comparing samples using sPCA is problematic for another reason: the requirement to choose a sign. Eigenvectors such as principal component vectors are unique only up to algebraic sign and the sign of a vector calculated by an eigensolver is arbitrary. Therefore, when required, a method or convention is needed to assign a sign or “orientation” deterministically. Historically in A/B compartment analysis, GC-content has served as a reference signal for choosing the orientation of the score vector, such that the positive phase has higher GC content than the negative phase. Without a similar universal reference signal that predicts higher-order vectors with high fidelity, no guidance exists for orienting sPC score vectors from different samples for comparison. Because joint PCA produces scores projected onto a common coordinate system, projections from different samples always share the same signs with respect to the basis vectors and therefore for the purpose of comparing scores the choice of sign no longer matters.

Taken together, our findings demonstrate that while sPCA may approximate global structure within individually analyzed samples, it fails to produce comparable components across samples. When comparing contact maps in detail across multiple biological conditions, a joint PCA approach is required. Overall, joint PCA provides robust, interpretable, and directly comparable embeddings across samples for comparative genome compartmentalization analysis.

### An atlas of long-range interaction profiles reveals distinct nuclear architectures associated with cell state

With a scalable and robust framework to embed long-range chromatin interaction profiles across biological contexts, we decided to apply *jointly-hic* to explore long-range autosomal interactions across diverse healthy human tissues and model organ development systems. We selected 89 high-quality Hi-C datasets from the ENCODE project, 4D Nucleome Consortium, and other published sources. These datasets include *ex vivo* mature human tissues, primary immune cells, and *in vitro* differentiation models simulating liver, heart, and pancreas development [41,42]. We applied stringent selection criteria for ENCODE datasets, requiring over 1 billion ligation pairs and more than 200 million non-negative pixels. Differentiation samples from the *in vitro* models were included irrespective of pair count. We generated jPCA embeddings of autosomal contacts for all samples at a 50-kb resolution, starting from *cis-*masked, *trans-*contact frequency matrices, producing 32 principal component vectors for each sample within a shared vector space (**Figure 2A, S.Figure 2A**). Heatmaps of the PC scores reveal contrasting profiles between samples, not only in PC1 but higher-order PCs as well, suggesting that higher-order PCs capture biosample-specific differences in genome architecture beyond traditional A/B compartmentalization.

**Figure 2.**
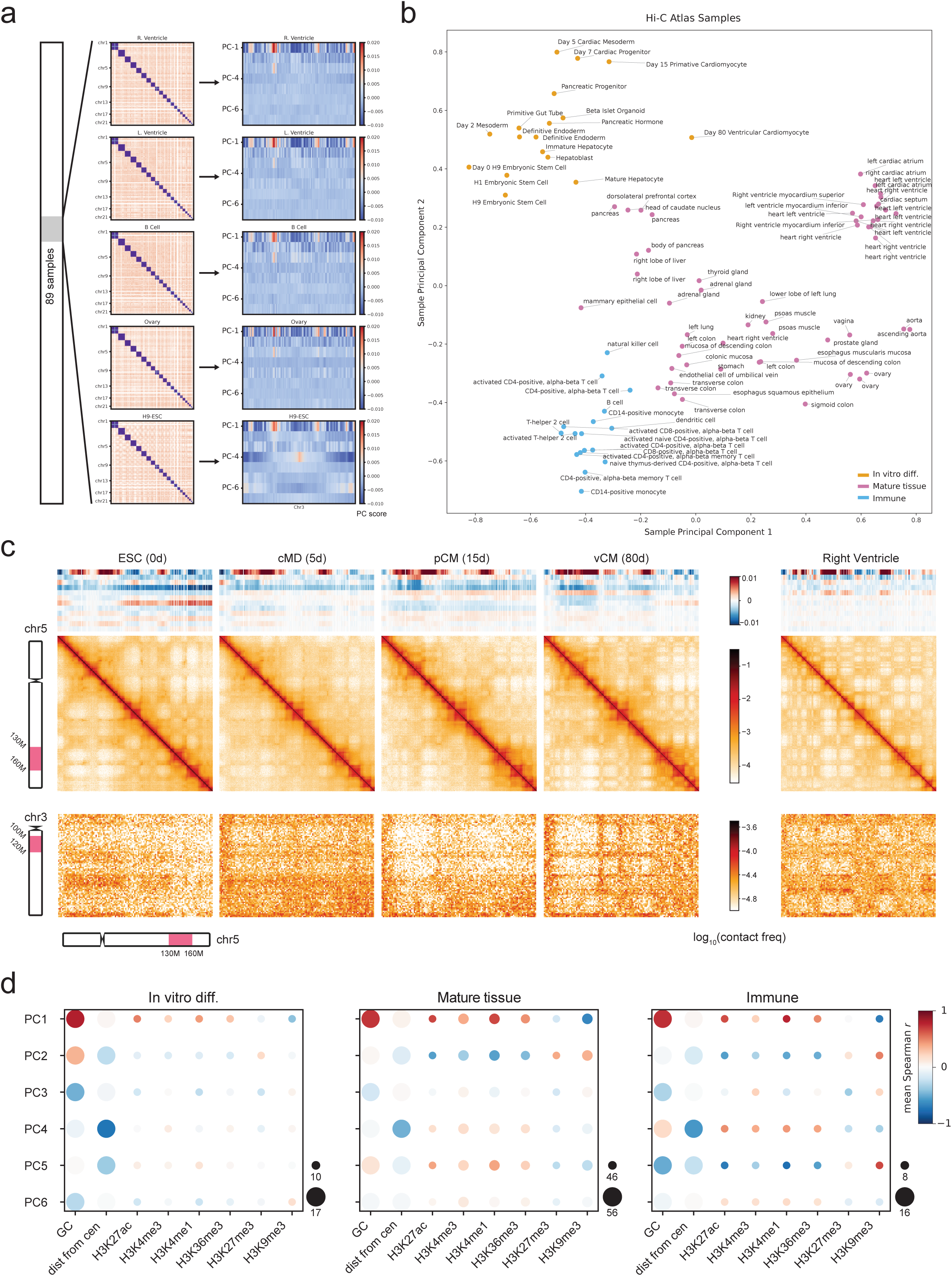
A locus-based atlas of long-range chromatin interaction profiles at 50-kb resolution across tissues reveals distinct genome architectural signatures between cell state categories. (A) Schematic of applying *jointly-hic* to 89 Hi-C datasets, yielding genome-wide principal components and PC scores at 50-kb resolution. Heatmaps of the first 7 PC score vectors for chromosome 3 shown as an example. (B) Overview plot of the sample similarity of long-range interactions via secondary PCA. Samples were projected onto the first two components of a secondary sample-level dimensionality reduction based on the first 10 PC score vectors from Hi-C. Points are labeled by biosample identifier and colored by cell state category. (C) Embeddings and contact frequency maps obtained from four stages (embryonic stem cell, cardiac mesoderm, primitive cardiomyocytes, and ventricular cardiomyocytes) along an *in vitro* cardiomyocyte differentiation trajectory (left) and a cardiomyocyte-rich *ex vivo* heart right ventricle sample (right). Top row: Heatmap of PC score vectors 1-12. Middle row: *cis* contact frequency maps showing a 30-Mb region on chromosome 5. Bottom row: *trans* contact frequency maps showing the same region against a 20-Mb region on chromosome 3. (D) Bubble plots depicting mean Spearman correlation coefficients between the first six principal component score vectors and biosample-matching genomic features for samples grouped into cell state categories *in vitro* differentiation (left), mature tissue (middle), and immune cells (right). The size of each bubble depicts the number of PC score vector + genomic track pairs contributing to the statistic.

To help interpret the biological relevance of our Hi-C-derived embeddings, for each sample we consolidated PC vector scores for all components across all genomic bins and conducted a secondary dimensionality reduction to compare the overall similarity of samples. This sample-level embedding clearly demonstrated proximity of similar tissues and cell types (**Figure 2B**). For example, cardiac tissues (heart ventricle, atrium, myocardium) cluster together, while immune cells (CD4+, CD8+) form another grouping. Moreover, hierarchical clustering of the sample-level interaction profiles grouped the samples into three high-level categories reflecting cell state: (i) H1 or H9 human embryonic stem cells and derived cells from *in vitro* differentiation experiments, (ii) mature immune cell samples, and (iii) other mature *ex vivo* adult tissue cell samples (**S.Figure 2B**).

Two additional mature samples grown in culture—mammary epithelial cells and umbilical vein endothelial cells—clustered with other *in vitro* samples, while localizing closer to immune samples in the sample overview PCA map, suggesting an effect of sample isolation conditions or sample clonality on long range interaction profiles (**Figure 2B**). To assess the impact of other potential technical and biological covariates, we examined batch effects and donor characteristics within ENCODE samples (**S.Figure 2C-F**). While laboratory and assay methodologies exhibited some separation, samples predominantly localized with other samples of similar tissue type. Donor age also displayed some gradient effects, but these often coincided with tissue types reflecting varying ease of sample acquisition (e.g., peripheral blood vs heart or brain).

Importantly, while *in vitro* differentiation samples were found to group closely together by similarity, in later stages of differentiation they gradually exhibit architectural characteristics reflective of their mature counterparts. For instance, in the sample-level embedding *in vitro-*derived mature hepatocytes aligned closely with human liver samples, ventricular cardiomyocytes cultured at 80 days converged toward heart ventricle samples, and endocrine pancreas samples drifted toward mature pancreas samples (**Figure 2B**). These progressions were evident in the original data by comparing, for example, both *cis* and *trans* contact frequency maps of the cultured cardiomyocytes to those of a mature right ventricle tissue sample (**Figure 2C**). We observed a trend toward the strong compartmental checkering pattern seen in the mature tissue sample, especially in the 80-day cultured sample.

To begin interrogating what type of information each component may be captured in our atlas, we grouped samples by cell state categories – *in vitro* differentiation, mature tissue, and immune – and for each category calculated the correlation of individual PCs with genomic variables and epigenomic features where matching data was available (**Figure 2D**). As expected, we find that PC1 scores consistently correlate with GC content across all samples and cell state categories, and exhibit consistent positive correlation with active histone marks (H3K27ac, H3K4me3, H3K36me3) and negative correlation with repressive or silencing ones (H3K27me3, H3K9me3). We find that across the three cell state categories, PC2 and PC3 scores exhibit inverse patterns of correlation with some features, including H3K27me3 signal. PC4 scores consistently exhibit a strong negative correlation with genomic distance from the centromere, suggesting that PC4 captures broad patterns of association between chromosome arms and centromere and telomere clustering in *trans* [13,21,22]. Intriguingly, the correlation profile of PC5 in the mature tissue category is the inverse of that in the immune category, while appearing weak across the board in the *in vitro* category, suggesting that PC5 scores are highly discriminative of the three categories of cell states. In the following section, we find evidence for other vectors that discriminate between cell state categories.

Collectively, our results suggest that broad long-range contact frequency signatures as encoded using our joint PCA approach reflect cellular identity and capture overall biospecimen similarity and functional differences between samples. Furthermore, our analysis at the sample level shows that joint PCA can discriminate between two categories of mature cells and between immature and mature cellular states, suggesting that samples within these cell state categories possess different baseline nuclear architectures.

### Long-range interactions distinguish broad chromatin states within and across cell state categories

To explore the collective interaction manifold more comprehensively at the individual locus level, we applied unsupervised clustering to the joint PC embeddings in two ways. First, we clustered the complete interaction profile of each individual locus restricted to its sample context of origin, thus taking the same genomic locus in two different samples as distinct observations. This approach, which we refer to as sample-locus (SL) clustering, finds a unified set of cluster assignments for sample-locus pairs, which may be interpreted in a similar way to subcompartment or IPG assignments in a single sample [21]. The second approach, which we refer to as ensemble-wide locus (EL) clustering, focuses on each locus’s complete set of interaction profiles across all samples in the atlas, thus considering each 50-kb genomic locus as a distinct entity. The latter approach is expected to group genomic loci together based on their sharing broadly coherent long-range interaction signatures across all of the samples in the atlas. Hence, while SL clustering can discern different interaction patterns of a given locus in different samples, EL clusters will consolidate interaction patterns that may differ from sample to sample but are correlated (e.g., loci having PC scores of opposite sign in two different samples). In both scenarios – SL and EL clustering – the final numbers of clusters, 11 and 8, respectively, were manually chosen by balancing of interpretability and model complexity.

To obtain a global overview of the two strategies, we visualized the results using a detailed locus-centric heatmap, where each column corresponds to a 50-kb genomic bin (locus) and each row corresponds to a genomic feature track, usually from a single sample (**Figure 3A**). The columns are sorted into eight EL locus clusters, labeled using Roman numerals. The EL clusters themselves are ordered by descending average GC content, and within each cluster the columns are ordered by ascending genomic distance from the centromere. The rows of the heatmap are organized into a series of blocks of feature categories. The first block depicts the frequencies of the eleven SL cluster labels for each locus as colored stacked bars. The subsequent groups of blocks arrange the PC score tracks from all 89 atlas samples. Scores corresponding to each PC rank appear in a unique block, and within each PC rank block, the rows are ordered according to the corresponding sample’s cell state category. The final sequence of blocks arrange 1,410 ChIP-seq signal tracks for various histone marks derived from biosamples matching those used in the atlas, with one separate block for each histone mark target, ordering the rows again by the sample of origin’s cell state category. Each row depicts the sample-wide z-score of ChIP-seq fold change over input signal for every 50-kb bin. To supplement the detailed heatmap with a visual aid, we also projected the full embedding of ∼44,000 50-kb loci from all 89 samples in the atlas using Uniform Manifold Approximation and Projection (UMAP) [45], and colored the sample-locus points in the projection by each of the major groupings in the heatmap: the corresponding sample’s cell state category (**Figure 3B**), the sample-locus’s SL cluster label (**Figure 3C**) and the corresponding locus’s EL cluster label across all samples (**Figure 3D**).

**Figure 3.**
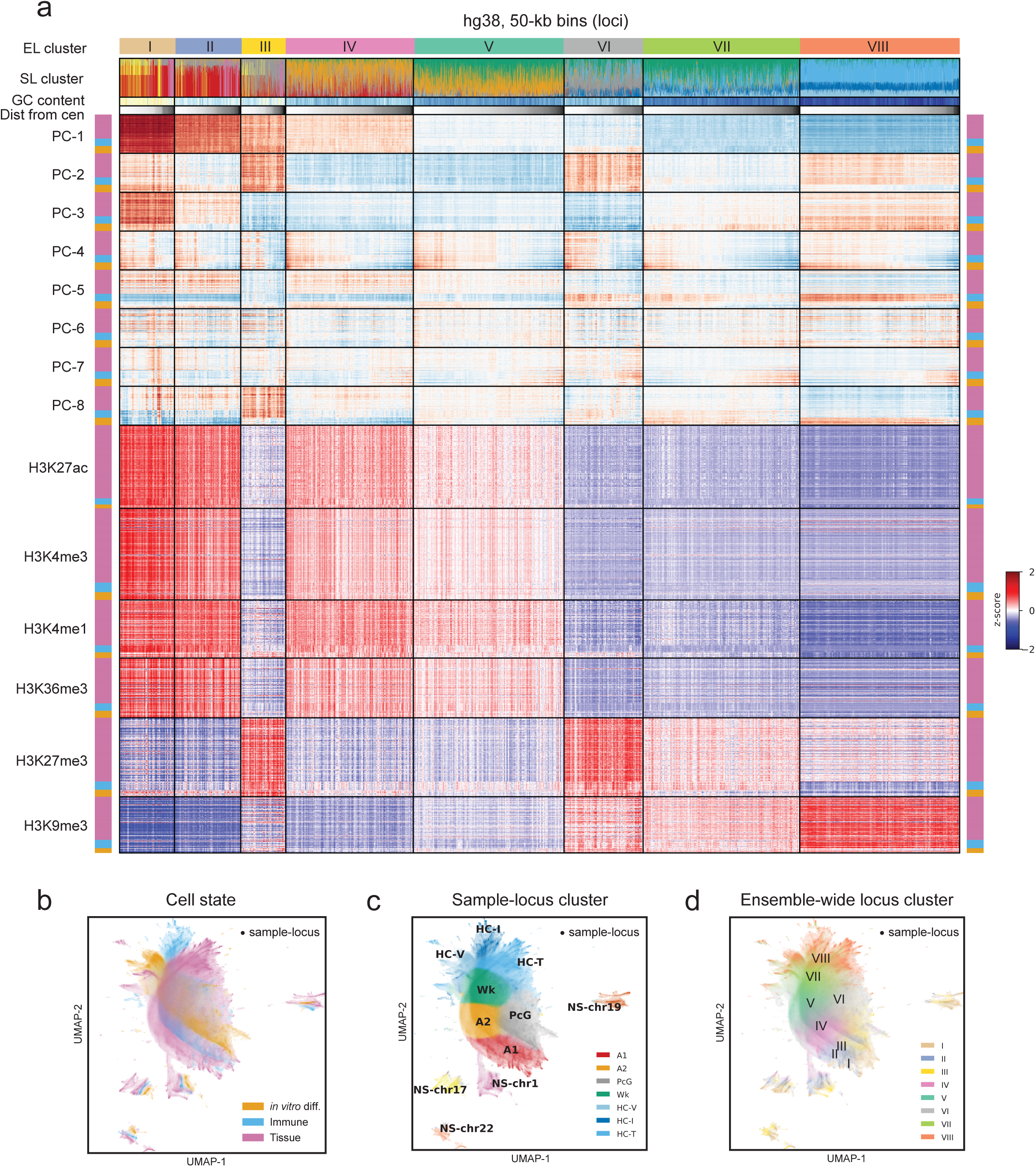
Locus-level cluster analysis and epigenetic characterization of the Hi-C interaction profile atlas. (A) Heatmap characterizing all 50-kb loci (genomic bins) mapped across 89 Hi-C samples. Columns correspond to loci and rows to different genomic tracks. Loci are grouped into ensemble-wide locus clusters by K-means (top labels in Roman numerals). Within each cluster, columns are ordered by genomic distance from the respective chromosome’s centromere. Second row: stacked bar charts of sample-locus cluster labels for each locus across 89 samples shown. Third row: GC content. Fourth row: genomic distance from the centromere. The next eight row blocks contain the top 8 PC scores for each locus from each of the 89 Hi-C samples, ordered vertically by cell state category (right/left colored bars). The last six row blocks contain z-score normalized ChIP-seq signals from ENCODE for biosamples matching those in the atlas, sorted vertically by cell state category (right/left colored bars). (B-D) Visualizations of UMAP embeddings derived from the full set of 32 PC scores for each sample-locus colored by (B) cell state category of the corresponding sample, (C) sample-locus cluster label, and (D) ensemble-wide locus cluster label of the corresponding genomic bin.

The detailed heatmap contextualizes the relationships between the Hi-C-derived principal component scores, SL and EL cluster assignments, chromatin state, and cell state category. For example, consistent with **Figure 2D**, PC4 displays a strong negative gradient with distance from the centromere within every EL cluster, supporting that PC4 captures a universal signature of interactions along chromosome arms in *trans*. This gradient appears steepest in *in vitro* samples, perhaps influenced by higher cell division rates, shorter gap phases, and resulting differences in cell cycle composition. By contrast, PC7 exhibits a centromere-telomere gradient in *in vitro* and immune samples but not in mature tissue samples, and the gradient appears more pronounced in clusters with lower transcriptional activity (low PC1 score). Components that discriminate cell state categories are also visible in the heatmap: PC2, PC3, and PC8 display unique signatures in *in vitro* samples relative to the others, while PC5 shows an inverted signature in immune samples relative to *in vitro* and mature tissue samples.

At the sample-locus level, consistent with the characterization of the traditional A compartment, most SL clusters with positive PC1 score are enriched for classical active chromatin marks such as H3K4me3, H3K27ac, H3K36me3, and H3K4me1 (**S.Figure 3A, 3C**). The SL clusters with the highest PC1 score were also enriched for POL2RA ChIP-seq and SON TSA-seq in all available cell lines where these marks were assayed, indicative of nuclear speckle-association (**S.Figure 3D, 3E**). SL clusters that exhibited strong enrichment for active marks without enrichment for repressive marks were annotated, in accordance with precedent, A1 (active, speckle-enriched) and A2 (other active) according to activity levels and SON enrichment. Additionally, of the top six SL clusters by mean PC1 score, four of them were small clusters harboring dense, speckle-associated regions unique to chromosomes 1, 17, 19 and 22, respectively (**S.Figure 3B**).

Surprisingly, among sample-locus pairs having the lowest average PC1 scores, we found three separate SL clusters of classical B-compartment sample-loci—which we labeled HC-V, HC-T, and HC-I—each almost exclusively associated with *in vitro* samples, mature tissue samples, and immune samples, respectively (**Figure 3B, 3C**). Interestingly, loci with positive PC2 scores and negative PC3 scores were associated with elevated H3K27me3 levels. Often these features co-occur with PC1 scores at similar levels to A2 bins, forming a separate SL cluster. We labeled this cluster PcG, as the features are indicative of polycomb-repressive states within broader regions of elevated transcriptional activity (**Figure 3A, S.Figure 3C**). Additionally we observed a SL cluster of loci lacking H3K27me3 which tended to exhibit marks of activity at levels lower than loci in A2 and/or marks of repression lower than those of the HC-V, HC-T, and HC-I clusters. We termed this latter cluster Wk for “weak”. The Wk loci are characterized by generally low histone modification signals for marks assayed by ENCODE but considerable variability depending on biosample context. To further examine regulatory activity across sample-loci in the atlas, we gathered ENCODE ATAC-seq data from mature tissue biosamples and estimated the most “active” regulatory elements as the top 100,000 (∼4%) of candidate *cis-*regulatory elements (cCREs) [35] per biosample by ATAC-seq signal. We found that A1 and the nuclear speckle SL clusters have the highest density of active regulatory elements, closely followed by A2. PcG and Wk EL clusters have low active cCRE density and HC-T is nearly depleted of active cCREs (**S.Figure 3F**).

At the ensemble-wide level, the EL clusters were found to be associated with epigenetic states that are broadly similar across the biosamples in the atlas with a few exceptions (**Figure 3A, 3D**). The first two EL clusters—I and II—are the most enriched for marks of transcriptional activity, encompassing most sample-loci with speckle-associated SL assignments. Clusters IV and V have progressively lower enrichments for active marks and comprise mostly sample-loci with the A2 and Wk SL cluster assignments. Interestingly, EL cluster VII consists of classical B-type loci that show only a weak enrichment for either of the two conventional heterochromatin marks (H3K27me3 or H3K9me3). Sample-loci with the Wk SL cluster assignment are primarily divided across between EL clusters V (weakly active) and VII (weakly repressive).

Interestingly, four EL clusters are strongly associated with the presence of conventional heterochromatin marks in a majority of samples (**Figure 3A-D**). EL clusters III and VI are enriched for H3K27me3, exhibit high PC2 and low PC3 scores in most samples, and comprise most sample-loci with the PcG assignment. Notably, EL clusters III and VI differ in their predominant PC1 status, with cluster-III loci corresponding to classic A-type loci (positive PC1) and cluster-VI loci corresponding to classic B-type loci (negative PC1). Together, this indicates that the presence of H3K27me3 is associated with different interaction signatures depending on the broader functional genomic context. EL cluster VII is depleted for active marks and exhibits mild levels of both H3K27me3 and H3K9me3. Finally, cluster VIII comprises the majority of sample-loci harboring the three cell state-specific SL inactive or heterochromatic cluster assignments, HC-V, HC-I and HC-T, and is strongly enriched for H3K9me3.

To explore the relationship between Hi-C embeddings and the proximity to nuclear landmarks, we gathered physical proximity-related data based on TSA-seq experiments in H1 hESC and HFFc6 cells [30,46] as well as Spatial Position Inference of the Nuclear Genome (SPIN) states derived from a combination of TSA-seq, DamID-seq and *cis* Hi-C data [47] (**S.Figure 4**). We found that projections along PC1 and PC2 were largely sufficient to separate H1 sample-loci by SPIN state, except for those labeled InteriorRepr1 from InteriorRepr2. This suggests that much of the information about spatial positioning is contained in the first two components of *trans* Hi-C data alone. As expected, we observed that Lamin B1 proximity was associated with the lowest PC1 scores, while SON proximity was associated with the highest PC1 scores, the A1 and nuclear speckle islands SL clusters, as well as with high PC3 scores. Centromere protein B (CENPB) proximity followed a similar pattern to SON, hinting at a preference of pericentric CENPB for the nuclear interior. On the other hand, nucleolus-associated proteins, NIFK and POLR1E, exhibited gradients that increased along PC2 towards higher levels of H3K27me3. These results suggest that while PC1 may follow a radial axis of spatial positioning in the nucleus, PC2 and PC3, interpreted spatially, may describe independent propinquities towards nucleoli (high PC2, low PC3) or nuclear speckles (low PC2 and high PC3). Together, these findings suggest that joint Hi-C embeddings contain signatures of both radial organization and the effects of nuclear body association.

**Figure 4.**
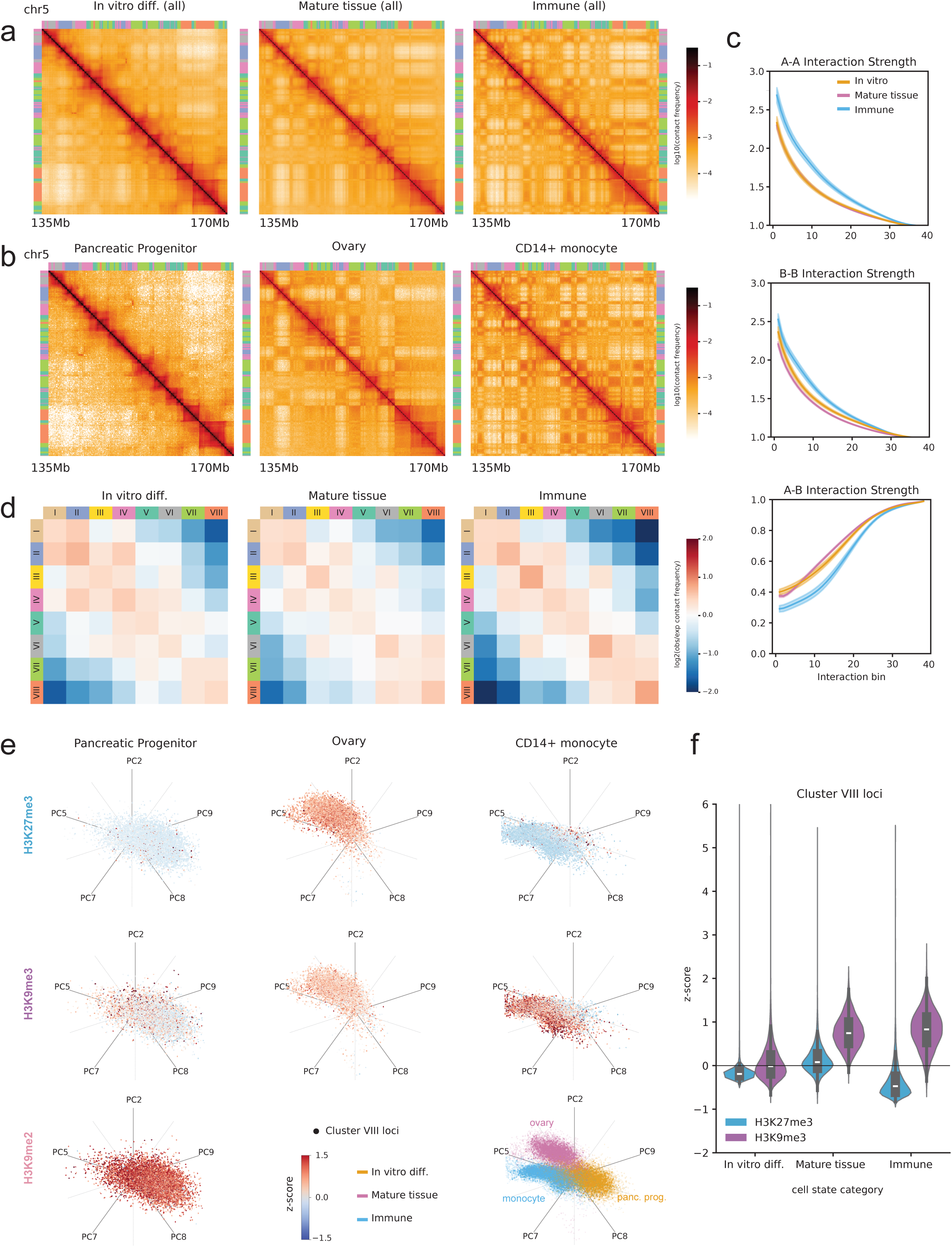
Samples in three cell state categories are associated with differences in the strength of heterochromatin compartmentalization and repressive histone mark composition. (A) Contact frequency maps generated from aggregating all samples within cell state categories show qualitative differences in average compartmentalization strength in *cis*. A 35-Mb region on chromosome 5 is displayed. (B) Contact frequency maps for representative samples from each cell state category: pancreatic progenitor cells (*in vitro* differentiation), ovary (mature tissue), CD14+ monocytes (immune cells). (C) Interaction strength plots derived from intrachromosomal saddle plots based on PC1 score quantile. Curves represent the mean interaction strength in each cell state category with standard error envelope. (D) Discrete saddle plots of average observed over expected contact frequency between loci in EL clusters across all samples in each cell state category. (E) Star coordinate scatter plots projecting genomic loci comprising EL cluster VIII along PC2, PC5, PC7, PC8 and PC9 for each of the representative samples from the three cell state categories. Points are colored by z-score of ChIP-seq signal for repressive histone marks H3K27me3 (top row) and H3K9me3 (middle row) from the same biosample. In the bottom left, the average H3K9me2 ChIP-seq signal from H9 ESCs and H9-derived mesoderm, cardiomyocytes, liver progenitors, and hepatocytes is overlaid [70]. In the bottom right, the point clouds from each of the three representative samples are plotted together and colored by cell state category. (F) Violin plots of H3K27me3 and H3K9me3 ChIP-seq z-scores across matching biosamples in each cell state category.

### Cell state-specific differences are associated with the heterochromatic composition of a common set of loci

It was surprising that the same large collection of genomic loci (EL cluster VIII) separated into three distinct transcriptionally inactive SL cluster assignments that are unique to each cell state category. Visually, these three sample-locus clusters diverged into three cell state-specific “lobes” in the UMAP embedding (**Figure 3C**). These observations prompted us to investigate whether compartmentalization differences in the predominantly silent portion of the genome may drive much of the global differences between the general architectures of these three categories of samples.

To observe the differences more directly, we aggregated all Hi-C datasets within each cell state category into pooled contact maps and visualized compartmentalization in patterns (**Figure 4A**). We found that the average immune cell map exhibited the strongest compartmentalization, characterized by prominent checkerboarding patterns, especially in EL cluster VII and VIII bins. The average mature tissue sample exhibited intermediate levels of compartmentalization of the same regions, with moderately defined checkering patterns, while *in vitro* differentiated samples displayed the weakest compartmentalization, characterized by smoother contact maps and less-defined genomic domain boundaries. We confirmed that these observations in the pooled maps were not averaging artifacts and were also evident in contact maps from individual samples from each category, such pancreatic progenitor cells (*in vitro*), ovary (mature tissue) and CD14-positive monocytes (immune) (**Figure 4B**).

To quantify these observations systematically, we generated saddle plot heatmaps of observed-over-expected contact frequency, consolidating genomic bins into 40 ranked groups based on their joint PC1 score quantile (**S.Figure 5A**). As observed in the pooled contact maps, the composite saddle plot of immune samples showed significantly higher classic A-to-A and B-to-B interactions and notably weaker interactions between A and B regions than *in vitro* and mature tissue samples in *cis*. Profiling the homophilic and heterophilic interaction strengths from the diagonals of individual samples’ saddle plots confirmed that immune cells demonstrated the strongest compartmentalization with higher self-affinity for bins with similar PC1 scores and greater segregation of between bins with opposing PC1 scores, while *in vitro* and mature tissues shared similar interaction strength profiles with respect to PC1 (**Figure 4C**). Additionally, we generated “discrete” saddle plot heatmaps of observed-over-expected contact frequency for each cell state category, grouping genomic bins into EL clusters as discrete categories (**Figure 4D, S.Figure 5B**). These heatmaps broadly agreed with the PC1 saddle plots. The inactive EL clusters VI, VII, and VIII were depleted for interactions with other EL clusters and the depletion was more severe in immune samples. Clusters VI and VIII both exhibited stronger self-affinity in immune samples as well. Interestingly, cluster VI was found to have lower affinity for clusters VII and VIII in *in vitro* samples than in mature tissue and immune ones.

**Figure 5.**
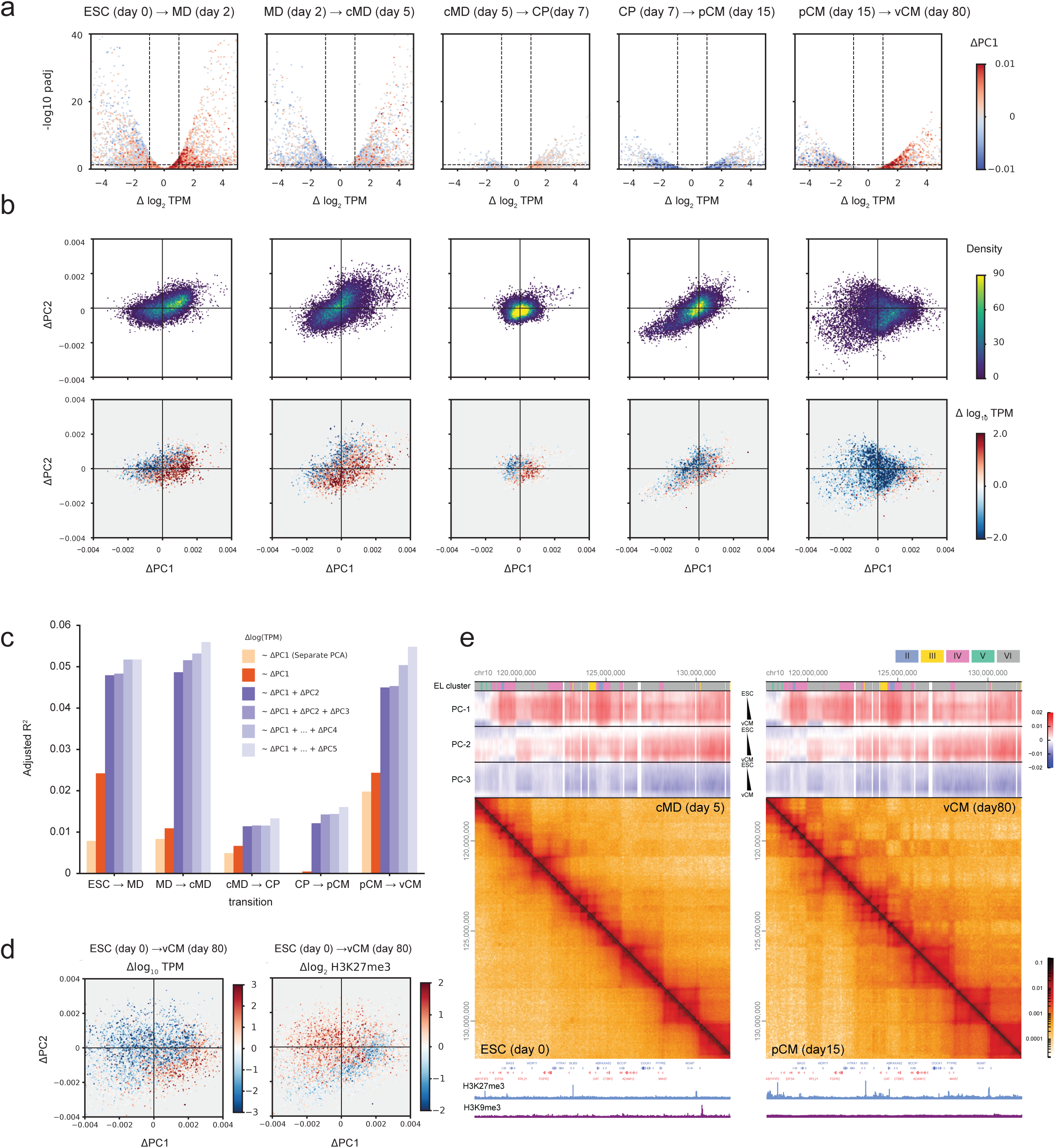
Atlas PC2 captures additional gene regulatory information during differentiation associated with downregulation of repressive marks within active regions and vice versa. (A) Volcano plots of differential gene expression between consecutive stages of cardiomyocyte differentiation, colored by change in PC1 score. (B) Scatter plots of change in PC1 score vs change in PC2 score for genes, colored by point density (top) and log fold change in gene expression (bottom). (C) Adjusted coefficients of determination for linear regressions predicting differential gene expression (log fold change in TPM) from differential PC scores for each stage transition. Models were trained on joint atlas PCs 1 through 5 cumulatively as well as for separately-calculated PC1. (D) Scatter plots of change in PC1 score vs change in PC2 score for genes between the first (ESC) and last stage (ventricular cardiomyocytes) colored by log fold chane in gene expression (left) and log fold change in H3K27me3 ChIP-seq signal between H9 cells and *in vitro* RUES-derived cardiomyocytes from ENCODE. (E) HiGlass visualization of contact maps from four stages (left view: ESC, cMD, right view: pCM, vCM) centered on a 15-Mb region of chromosome 10. EL cluster labels and PC score tracks for all six differentiation stages are displayed on top of both views. Underneath both views are gene annotations and BigWig tracks of H3K27me3 and H3K9me3 from H9 (left view) and RUES-derived cardiomyocytes (right view).

Comparing these results to the depiction of EL clusters VII and VII in **Figure 3A**, we noticed differences in enrichment of marks H3K27me3 and H3K9me3 between cell state categories. In the case of EL cluster VII, H3K9me3 enrichment is relatively consistent across samples. However, while H3K27me3 is generally—albeit weakly—enriched in these loci for both tissue and immune samples, the same mark is generally depleted in the same loci in *in vitro* samples. In the case of cluster VIII, loci are enriched for H3K27me3 in mature tissue samples, but depleted for the same mark in both immune and *in vitro* samples. Simultaneously, a strong enrichment for H3K9me3 is seen in tissue and immune samples yet not in *in vitro* samples. Given that PC basis vectors 2, 5, 7, 8 and 9 appeared to distinguish sample-loci by cell state category, we projected all the genomic loci in cluster VIII from characteristic samples from each category—pancreatic progenitor, ovary, and CD14+ monocyte—onto a multivariate star coordinates plot [48] (**Figure 4E**). Coloring the points by sample confirmed a clear separation by cell state category, as expected. Splitting the points from each sample into separate plots, we then colored them by histone modification z-scores for H3K27me3 and H3K9me3, which supported the observation that cluster VIII loci have a mild to neutral enrichment of H3K27me3 in tissue samples, while being depleted for H3K27me3 in *in vitro* and immune samples. At the same time, these same loci are strongly enriched for H3K9me3 in tissue and immune samples but not in *in vitro* samples. Finally, the observed asymmetries in H3K27me3 and H3K9me3 levels in cluster VIII loci were confirmed by plotting distributions of sample-loci z-scores across the three cell state categories (**Figure 4F**).

Interestingly, we found that for *in vitro* samples, genome-wide, broad H3K9me3 enrichment is largely limited to peri-centromeric, telomeric regions and characteristic domains on chromosome 19, while most of the peripheral, lamin-associated regions in the genome are enriched for H3K9me2 rather than H3K9me3 (**S.Figure 5C**). Together, these results suggest that chromosome organization is influenced not by simple binary affinities between heterochromatic regions but are affected by combinatorial differences in the epigenetic composition of heterochromatic or pre-heterochromatic domains.

### Changes in jointly-derived PC scores coherently predict gene expression dynamics during *in vitro* development

Previous work has associated compartmentalization changes, as transitions between B compartment (negative PC1 scores) and A compartment (positive PC1 scores), with transcriptional activity shifts during cellular differentiation [49,50]. To systematically investigate this relationship using our atlas, we turned again to the comprehensive six-stage *in vitro* cardiomyocyte differentiation dataset comprising matched Hi-C, RNA-seq, and ATAC-seq assays [42] and performed differential expression analysis between successive development stages. We colored volcano plots of differential gene expression by changes in jPC1, which revealed that genes with high expression changes tended to have high changes in jPC1 scores (**Figure 5A**), consistent with the hypothesis that classic A/B compartmental status changes contribute to gene expression changes during development. However, the extent to which higher order principal components capture meaningful expression and regulatory changes remained unclear.

To determine whether higher-order PCs capture additional layers of regulatory information, we examined the relationship between changes in jPCA scores (ΔjPC) and changes in gene expression. Scatter plots of gene-level ΔjPC1 versus ΔjPC2, colored by log_10_ TPM fold change, showed clear diagonal gradients, indicating that a combination of ΔjPC1 and ΔjPC2 better explains transcriptional dynamics than ΔjPC1 alone (**Figure 5B**). A similar trend was observed when coloring by changes in chromatin accessibility at ENCODE cCREs (**S. Figure 6A, 6B**). To quantify this information and examine the contribution of each PC to gene expression dynamics, we fit a series of linear regression models using ΔjPC1 through ΔjPC5 to predict log TPM fold change across each stage transition (**Figure 5C**). As a baseline, we also included models using ΔsPC1 from separately-calculated PCA on each stage. Across all transitions, ΔjPC1 consistently outperformed ΔsPC1, as measured by adjusted R², suggesting that the joint embedding not only provides a more biologically coherent coordinate system for comparative analysis, but one that better captures transcriptional changes in a dynamic experiment. Importantly, the inclusion of ΔjPC2 substantially improved model performance, explaining up to 5% of the variance in log TPM changes. We observed that performance increased only marginally beyond ΔjPC2, indicating that most of the predictive signal is captured by changes in the first two joint components.

The observation that changes in higher order components capture information relevant to gene expression dynamics independent of ΔPC1 is supported by our earlier observation that a pattern of PC2 and PC3 in the atlas are associated with the two EL clusters enriched for the repressive histone modification H3K27me3 across most samples (**Figure 3A**). Namely, we observe that PC2 and PC3 take on anticorrelated positive and negative signs, respectively, in EL clusters III (H3K27me3-enriched with high PC1 score) and VI (H3K27me3-enriched with low PC1 score), while taking on similar values in the other quiescent EL clusters VII and VIII. This suggested that the combination of PC2 and PC3 can discriminate between different types of silencing chromatin states, such as Polycomb-repressed states within peripheral, gene-poor or interior, speckle-rich regions, and that the inclusion of PC2 and to some extent PC3 captures higher-order patterns of chromatin silencing than is possible using the traditional A/B compartment score. To test this, we collected H3K27me3 ChIP-seq data from ENCODE for H9 cells (the first stage) and for a different *in vitro* cardiomyocyte sample derived from RUES cells, as a proxy for the last stage. Coloring gene-level ΔjPC1 and ΔjPC2 for ES cells versus ventricular cardiomyocytes by log fold change in TPM and H3K27me3 showed inverted gradients along the same direction, where lower gene expression and increased H3K27me3 are associated with decreases in both jPC1 and jPC2 and vice versa (**Figure 5D**). Supporting this, we could find examples of large regions of positive PC1 (classic A) and positive PC2 score in early differentiation stages, that fragment in later stages, forming interspersed troughs in PC2 accompanied by the accumulation of H3K27me3 as well as stronger compartmental checkering (**Figure 5E**). Together, our results show that jointly-derived higher-order components from Hi-C data can provide significant information about long-range interactions relevant to gene expression and epigenetic state dynamics beyond the classic compartment flipping model.

## Discussion

Traditional methods for the analysis of long-range contact frequencies have substantial limitations for integrative analysis because separately applied matrix decomposition methods produce low-dimensional embeddings that are projected onto dissimilar basis vectors. To address this need, we developed *jointly-hic* to normalize and apply PCA to multiple contact frequency maps simultaneously, which projects each dataset onto a common set of basis vectors, allowing for much deeper comparative analyses than previously accessible. Importantly, the *jointly-hic* toolkit implements this methodology while meeting the computational constraints for large-scale integrative studies.

In this study, we demonstrate that joint linear embeddings capture coherent and biologically meaningful features, and that these features provide superior and consistent interpretations across the stages of an *in vitro* cardiomyocyte differentiation model system. Further, we show that joint PCA decompositions are robust to sequencing depth variations and background random ligation noise. A key result is that higher-order jointly-calculated PC score vectors (i) capture meaningful information beyond the traditional A/B compartment score and (ii) are consistent and comparable across samples, which is not the case for separately calculated embeddings. While joint PC1 score vectors align quite closely to the traditional A/B compartment vector, we showed that higher-order PCs capture important orthogonal information, including patterns related to chromosome arm alignment and diverse repressed or heterochromatic states. Understanding the mechanistic origins of these higher-order features will help unravel how various cellular and molecular processes and chromatin states influence genome architecture and how such changes drive the evolution of cell state during development.

Signatures of distinct nuclear architectures among broad cell state categories emerged along several components in our atlas-wide joint embeddings. While the influences of nuclear morphology and sample heterogeneity likely play a role, we found that immune cells, mature tissues, and *in vitro* ESCs and ESC-derived samples were associated with differences in the epigenetic composition and apparent maturation state of heterochromatin. Immune cells exhibited H3K9me3 with very low levels of H3K27me3 in heterochromatic loci, whereas mature tissues showed a H3K9me3 predominant with mild signal levels of H3K27me3. In contrast, both ESCs and ESC-derived cell types *in vitro* contained putatively “immature” heterochromatin states characterized by H3K9me2 with minimal presence of the trimethyl mark. This observation, which may be linked to common human embryonic stem cell lines being in a primed, epiblast-like state [51,52], has direct consequences on genome compartmentalization.

We further identified an intriguing ensemble-wide cluster of loci (EL cluster VII) exhibiting a preponderance of weakly repressive chromatin marks that did not conform neatly to classical heterochromatin definitions, raising the possibility of alternative or transitional heterochromatin states. While EL cluster VI appeared marked primarily by H3K27me3, suggestive of facultative heterochromatin, and VIII was predominantly marked by H3K9me3 in mature cell types, suggestive of constitutive heterochromatin, EL cluster VII contains a mixture of the two marks. One possibility is that these regions represent bivalent states marked by both H3K27me3 and H3K9me3, consistent with recent observations from K9 methyltransferase knockout models, which suggest competitive interplay between these histone modifications [53]. Another possibility is that there is a stronger likelihood of cross-reactivity between ChIP-seq antibodies for their trimethyl target in these lower signal regimes. Further validation, including perturbation experiments of chromatin modifiers, would be invaluable in clarifying the nature of these chromatin states.

One caveat of the *jointly-hic* approach, as an unsupervised method, is that the jointly calculated components are dependent on the input data. For example when computing joint PCA on only the cardiomyocyte stages, basis vector PC2 correlates with centromere-telomere distance, but when computing joint PCA on the entire atlas of 89 samples, PC4 captures this feature instead. In the atlas-wide embedding, we showed that changes in PC1 and PC2 carry information that is independently predictive of differential gene expression. The differences are likely due to the fact that “Rabl-like” centromere-to-telomere contact frequency patterns in *trans* are more salient in the *in vitro* differentiation samples, and less so in the post-mitotic and more heterogeneous *ex vivo* tissue samples which make up a large fraction of the atlas. Therefore, in general, when an unsupervised joint PCA is performed in any given study, the biological interpretation of the resulting principal components will need to be determined for that work.

Alternatively, a very large and comprehensive atlas of biosamples could be used to produce a well characterized set of basis vectors that could then be reused by the community at large. New samples can be projected into such a reference latent space by a simple linear transformation. Furthermore, the incremental PCA algorithm used by *jointly-hic* is iterative and stateful, such that new samples may be inexpensively added to an existing atlas of training examples to continually refine the consensus basis vectors [54]. By linear projection, joint PCs produce parametric and biologically meaningful embeddings, which can support a broad range of downstream analyses. For example, one promising avenue would be using joint PCA embeddings of Hi-C data as harmonized features for training emerging sequence-based deep learning models [55–60], thereby incorporating global long-range 3D genomic data into existing frameworks .

The current study has limitations. While our incremental PCA method is not intrinsically limited by genomic resolution, due to the sparse coverage in *trans* even for some of the most deeply sequenced Hi-C datasets we limited our analyses to a 50-kb bin size, which misses finer compartmental structure [16,61,62]. Our reliance on exclusively interchromosomal contact frequency data and a single genomic resolution limits the amount of information gleaned about chromosome organization since *trans* interactions are much less frequent than those occurring in *cis* and therefore also more sensitive to noise. However, the approach does offer advantages, including the ability to obtain genome-wide embeddings that are automatically harmonized not only across samples but also across chromosomes, and avoiding the need to account for polymeric distance dependences. Focusing on interchromosomal contacts also significantly decouples patterns of compartmentalization and long-range interactions from the patterns of local regulatory interactions and loop extrusion-driven TAD-scale dynamics, even though the latter does exert influence on the former. Future work extending our method to *cis* data through chromosome-specific joint decomposition will provide even more informative and higher resolution embeddings which may enhance the sensitivity of case/control studies and perturbation experiments. For example, existing methodologies developed for the differential analysis of the traditional A/B compartment vector may be extended to take higher-order jointly-derived PCs into account [38]. Another potential application of incremental PCA for further development is in the analysis of compartmentalization in single cell Hi-C [63].

In conclusion, our work provides an approach for integrative analysis of genome compartmentalization and patterns of long-range interactions across diverse biological contexts in a manner that is robust, scalable, and biologically informative. Our findings establish that, when learned jointly, higher-order latent components can be used to provide information about genome organization and biological information reflective of cellular identity, chromatin state, and developmental trajectories. By enabling direct comparisons across Hi-C datasets, *jointly-hic* advances our capacity to interpret 3D genome dynamics, laying the groundwork for future integrative studies in development and disease.

## Methods

### Joint principal component analysis

The traditional matrix-based approaches to the dimensionality reduction of Hi-C data correspond closely or exactly to PCA and the differences between methods lies largely in the pre-processing of the input matrix. All can be cast as the spectral (eigen) decomposition of a normalized or “observed-over-expected” contact matrix following mean-centering [13] or calculating its correlation matrix [12]. If the final input matrix *X* ∈ ℝ^*n* × *n*^, where *n* is the number of genomic bins after coverage-based filtering, is symmetric (*X^T^* = X) and balanced (constant row/column sums), the former approach produces the same vectors as a conventional PCA, up to a scale factor, which can be expressed as the eigenvectors of the covariance matrix 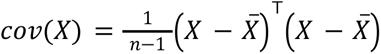. The latter approach corresponds to a correlation-based PCA, which is equivalent to applying an additional z-score transformation to the features of *X* before performing conventional PCA. In general, both types of PCA are normally computed by performing a singular value decomposition (SVD) of the mean-centered final input matrix, where right singular vectors correspond to principal components.

To perform a joint PCA factorization on an arbitrary number of contact frequency profiles (observations) over many input contact matrices, *jointly-hic* uses the IncrementalPCA estimator from *scikit-learn* [40], which performs an iteratively-updated SVD in mini batches, returning the most significant singular vectors [19,54]. This algorithm has constant memory complexity, on the order of the batch size times the number of features. Altogether, the *jointly-hic* pipeline consists of the following steps:

1. Genome-wide balanced contact frequency matrices are loaded from multiresolution cooler [64] (mcool) files at 50-kb resolution. Only autosomes (chromosomes 1-22) were included for the analyses in this study.
2. Intrachromosomal contacts (*cis* regions) are removed and replaced with randomly sampled *trans* pixels from the same row/column, as in [65].
3. Blowout pixels, defined as above the 99.5th percentile threshold, were clipped. Pixels below the first percentile were set to 0 to remove additional poor coverage bins.
4. Contact matrices are rebalanced so that all rows and columns sum to 1.
5. Genomic bins that get masked in any of the input samples are combined into a union list and pre-processed input matrices are saved to disk for further stages.
6. For each pre-processed input matrix, incremental PCA is run, updating in chunks of 10,000 rows per iteration via the ‘partial_fit’ method. After each input matrix is exhausted, the next is loaded.
7. Once the final joint PCA model is fit, a second pass through the data is performed to transform the input matrices into PC score vectors (projections).

In general it is challenging to determine purely technical batch effects to correct for *a priori*. However, we noticed that increased background ligation noise in our simulations tended to decrease the overall dispersion of the sample-loci from the origin for a given sample isotropically. To correct for such sample-to-sample differences in background noise levels, we divide the matrix of PC score vectors derived from each sample by the Frobenius (matrix) norm for that sample, which normalizes how “spread out” each sample’s point cloud is around the origin. The final PC score vectors are rescaled to the global norm of the original projections.

### HCT116 downsampling and noise injection

We verified the reproducibility of embeddings from *jointly-hic* using the unsynchronized, untreated RAD21-Auxin-inducible degron (AID) HCT116 dataset from [43]. The pseudodiploid colorectal cancer cell line was processed using *distiller-nf*, and subjected to downsampling and synthetic injection of random ligations. Non-nuclear, sex chromosomes, and three large autosomal translocations were excluded as previously described [21]. Downsampling was performed using *cooltools sample* [15]. The number of removed counts was replaced with randomly generated ligation pairs drawn uniformly across the genome to restore total read counts. Using *jointly-hic*, we computed joint PCA embeddings for the original, downsampled, and downsampled with artificial noise samples.

### Cardiomyocyte differentiation separate and joint PCA

We applied PCA separately to each contact matrix, generating sPCA latent embeddings for each sample. Independently, we also applied the full *jointly-hic* processing pipeline to generate jPCA embeddings. Genomic features including GC content and centromere-telomere distance computed at the same loci were merged with the sPCA and jPCA embeddings with the help of *bioframe* v0.8.0 [65]. Due to the arbitrary algebraic sign of sPC score vectors from stage to stage, we calculated absolute values of Pearson correlation coefficients between all sPC and jPC score vectors within and across stages. Pairwise angles between sPC2 and sPC3 score vectors in different differentiation stages was computed by calculating cosine similarities and converting to the corresponding acute angle in degrees.

### Biosample atlas data curation

We curated an atlas comprising 90 human Hi-C datasets. We included 73 Hi-C samples from the ENCODE data portal, containing greater than 1 billion ligation pairs and greater than 200 million non-negative bins, from *ex vivo* and primary culture biosamples. We excluded immortalized and cancer cell lines and samples with abnormal karyotype or structural variants. We also included human *in vitro* differentiation model systems representing the heart [42], pancreas [41] and liver (4DN consortium) from published studies and the 4DN consortium. After applying joint PCA, we excluded one sample (ENCODE accession ENCSR797MWY, aorta tissue from a 41 year old female) from all further analysis. Its latent embeddings differed substantially from all others, and upon visual inspection of the contact map, this sample showed pervasive and highly unusual chromosome-level fluctuations in contact frequency in *trans*, suggestive of extensive aneuploidy or some perhaps some other phenomenon, such as homologous chromosome pairing recently reported in adult aortic endothelial cells [66]. Sample metadata including experiment and file accessions and biosample information are available in Supplemental Table 1. In total, we curated an atlas of 90 Hi-C datasets, and kept 89 for all analysis in this study.

We prepared Hi-C datasets in mcool files for analysis, visualization and processing using *jointly-hic.* ENCODE samples were downloaded from the ENCODE portal as hic format files. For these samples, read quality-control, alignment to the hg38 genome, filtering, and conversion to contact frequency matrices were already performed with the ENCODE implementation (github.com/ENCODE-DCC/hic-pipeline) of the Juicer pipeline [67]. The hic format files were converted to the cooler format using *hictk* (version 0.0.10) at 1000 base pair resolution [68]. We used *cooler [64]* to “zoomify” and re-balance these to multi-resolution cooler (mcool) files. Sequencing read data from differentiation models were downloaded as FASTQ files from Gene Expression Omnibus with the accession GSE116862 for the cardiac and GSE210524 for the pancreas systems. Data for the hepatocyte differentiation model was obtained directly from collaborators and is available via the 4DN data portal under accessions provided in Supplemental Table 1. Raw sequencing reads were processed using *distiller-nf* (version 0.3.4) through read quality-control, alignment to the hg38 genome, filtering, and conversion to contact frequency matrices (github.com/open2c/distiller-nf). To complement the atlas of Hi-C datasets within a broader functional context, we curated collections of histone ChIP-seq, transcription factor (TF) ChIP-seq, and ATAC-seq. We queried ENCODE for all matching files, choosing data processed by ENCODE4, aligned to hg38, with the status “released,” and a biosample name matching one of the Hi-C atlas biosamples. In total, we collected 1,410 Histone ChIP-seq tracks (Supplemental Table 2), 321 TF ChIP-seq tracks (Supplemental Table 3), and 185 ATAC-seq tracks (Supplemental Table 4). Each track was downloaded from the ENCODE portal in BigWig format [69]. We integrated spatial landmark data from the 4DN data portal for DamID-seq targeting AP3D1 and LMNB1, and TSA-seq targeting LMNB1, SON, POLR1E, NIFK, and CENPB. The tracks and accessions are available in Supplemental Table 5. We downloaded processed BigWig files for LMNB1 ChIP-seq and H3K9me2 ChIP-seq from GEO, with the accessions in Supplemental Table 6 [70]. Genome browser visualizations were generated using HiGlass [71] and *higlass-python* (github.com/higlass/higlass-python).

### Embedding vectors and signal track aggregation

Following joint PCA, metadata tables were generated containing experiment metadata about biosample names, accessions, and mappings between file names and experiments in YAML format. Using *jointly-hic*, we created a custom HDF5-based database called a “JointDB” containing all jPCA embeddings from Hi-C at 50-kb resolution and corresponding epigenetic signal data from BigWig files of ChIP-seq, ATAC-seq, and other epigenetic profiling assays aggregated at the same resolution, as well as experiment metadata from all included samples and tracks. The JointDB can be queried as pandas dataframes using the *jointly-hic* software.

For each cell state category, we collated 50kb bin-level PC scores with matching signal tracks from the same cell state category. These tracks included GC content and genomic distance from centromere. We calculated Spearman correlation coefficients of the 50-kb signal tracks separately for each sample PC score vector by each biosample-matching matching track, and displayed the mean correlation coefficient in bubble plots sized by the number of tracks contributing to the calculation. We also collated all 50kb bin-level ChIP-seq profiles of H3K27me3 and H3K9me3 for each cell state category, mean-centered and scaled them by standard deviation, and generated violin plots to show the scaled signal distribution among the three categories.

### Atlas sample-level overview

We generated an overview of the Hi-C atlas by analyzing the full sets of genomic bin-level PC scores for each sample. We concatenated the first 4-32 PC score vectors from the *jointly-hic* embeddings for each of the 89 biosamples and applied a secondary PCA with two output components to visualize sample-level similarities. These embedding plots remained visually stable after including 5 or more top PC score vectors. We also applied hierarchical Ward clustering to the sample-level flattened feature vectors consisting of the first 12 PC score vectors and computed the euclidean distance map between them using *scipy* [72] and *fastcluster* [73].

### Atlas locus and sample-locus cluster analysis

For sample-locus clustering, we applied Leiden clustering (leidenalg v.0.10.2) to all sample-locus embeddings simultaneously over a range of resolutions including 0.1, 0.2, 0.3, 0.5, 0.8 and 1.0 and using 500 as the nearest neighbor cutoff [74]. For ensemble-wide locus clustering, we applied K-means clustering using *scikit-learn* to the locus-level embeddings, i.e. to the full set of sample-level embeddings for each locus, where each input vector corresponds to the concatenation of 32 PC scores from each sample. K-means was computed for a range of k including 5, 6, 7, 8, 9, 10, 12, 15, 20. UMAP (umap-learn v0.5.6) [75] visualizations of the sample-locus PC projections were computed for a range of nearest neighbors of 30, 100 or 500.

### Cardiomyocyte differentiation RNA-seq analysis

FASTQ files of RNA-seq data from six time points, each represented by two replicates: day 0, day 2, day 5, day 7, day 15, and day 80, corresponding to the Hi-C data, were downloaded from SRA. We used Salmon v1.10.3 for transcript-level quantification against the Gencode v46 transcripts annotations and GRCh38 primary assembly with the “validateMappings” and “gcBias” flags [76]. Then we used tximport v.1.30.0 to aggregate transcript abundances to the gene level [77]. A sample metadata table was generated to map each sample to its corresponding time point and experimental stage, which enabled a time-based design in the subsequent differential expression analysis. We applied PyDESeq2 v0.5.1 to compute differential gene expression statistics between consecutive stages [78]. We overlapped the genomic coordinates of increasing and decreasing differentially expressed gene TSSs with atlas bins and their jPCA scores at all developmental stages with *bioframe*. For each sequential stage transition, we used *statsmodels* [79] to fit linear models to predict log fold changes in RNA-seq TPM using differences in PC scores as regressors.

### Cardiomyocyte differentiation ATAC-seq analysis

For the cardiomyocyte time-course experiments, we downloaded ATAC-seq sequence data for the six developmental stages, each with two replicates, corresponding to the RNA-seq and Hi-C experiments. We processed the data using the ENCODE ATAC-seq pipeline (github.com/ENCODE-DCC/atac-seq-pipeline). We generated signal BigWigs directly from the filtered alignment files. To correct for transposase cutting biases, we first used the alignmentSieve utility (part of the deepTools suite, version 3.5.5) with the “ATACshift” parameter to adjust each BAM file [80]. The shifted BAM files were then sorted and indexed using samtools v1.13 [81]. Following sorting, coverage tracks were created with bamCoverage (deepTools) using a 1 bp bin size, with normalized coverage by the read-per-genome-coverage (RPGC) method, exact scaling, and excluding duplicate reads. Each coverage track was stored in BigWig format, and also averaged over 50-kb tiling genomic bins and added to the JointDB database. We quantified ATAC-seq coverage at ENCODE candidate cis-regulatory elements (cCREs, version 4 [35]) through a read count based approach. First, cCRE regions were defined in a featureCounts SAF (Simple Annotation Format) file containing genomic coordinates. We then used featureCounts (v1.5.3) in pair-end mode to count reads overlapping the cCRE intervals across all replicate BAM files [82]. We applied PyDESeq2 v0.5.1 to compute differential accessibility statistics between consecutive stages [78].

### Cell state category contact maps and saddle plots

To visualize differences between the cell state category groups, we created “mega-maps” by merging all relevant contact frequency matrices using the *cooler merge* command, followed by *cooler zoomify* to generate multi-resolution, balanced matrices. These aggregated contact maps capture an averaged contact pattern representative of each class. Using *cooltools*, we calculated per-chromosome expected contact frequencies (*cis* and *trans*) as input for saddle plot generation. The saddle function in cooltools was then applied to calculate saddle plot matrices from each individual sample contact matrix. Continuous saddle strength metrics were calculated from each saddle plot matrix as previously described, providing a ratio of intra-compartment interactions to inter-compartment interactions over successive quantile bins along the eigenvector. The average and standard error envelope of these curves for each cell state category were plotted. A slightly modified version of the *cooltools* saddle function was used to calculate “discrete” saddle plots over EL cluster categories.

### Locus-level heatmap, embedding plots, and overlays

The locus-level heatmap was generated with matplotlib from 50-kb PC score vectors and signal tracks from the JointDB using custom scripts. Rows representing score vectors and signal tracks were annotated grouped based on cell state category and ordered within each group. Columns representing 50-kb bins were grouped by EL cluster assignment. The EL clusters themselves were ordered and numbered by mean GC content, and within each group the columns were ordered by genomic distance from the centromere.

To avoid issues with overplotting of very dense scatter plots and support the overlaying of various quantitative and categorical variables associated with sample-loci in our various projection visualizations, we used the the *matplotlib* [83] extension of the *datashader* package [84] (datashader.mpl_ext.dsshow). This tool allowed for accurate rendering of two-dimensional point density plots and other aggregates of our PCA and UMAP sample-locus embeddings as well as gene-level scatter plots and volcano plots. With dsshow, point counts or alternate aggregations over points, such as mean values of a signal track, are generated over pre-defined grids in 2D cartesian space and rendered as raster images with a user-defined color map. Categorical aggregations are performed by color-encoding the categorical labels of points, quantitatively scaling the aggregates for each category using transparency, and compositing the resulting color channels. We also implemented a plotting tool to generate radial “star coordinates” plots from a given joint PC subspace using matplotlib’s scatter function for vector-based rendering or dsshow for raster-based rendering (https://gist.github.com/nvictus/f90b32503a3da126e2c72fda13a7119c). These plots project n-dimensional data points onto a two-dimensional circle with each dimension axis separated by equal angles around the origin.

## Data and Code Availability

The *jointly-hic* software package is open source and available at github.com/abdenlab/jointly-hic. The datasets analyzed in this study are listed in supplemental tables 1-6 and available from ENCODE, 4DN, or GEO under the corresponding accessions.

## Acknowledgements

We thank Guoyun Chen for contributions to *jointly-hic*. We thank Aleksandra Galitsyna, Greg Andrews, Félix Raimundo, Johan Gibcus, Zeeshan Siddiqui, members of the Mirny, Dekker, Maehr labs and the 4DN Center for 3D Structure and Physics of the Genome, and members of Open2C for feedback and helpful discussions.

---

**S.Figure 1.**
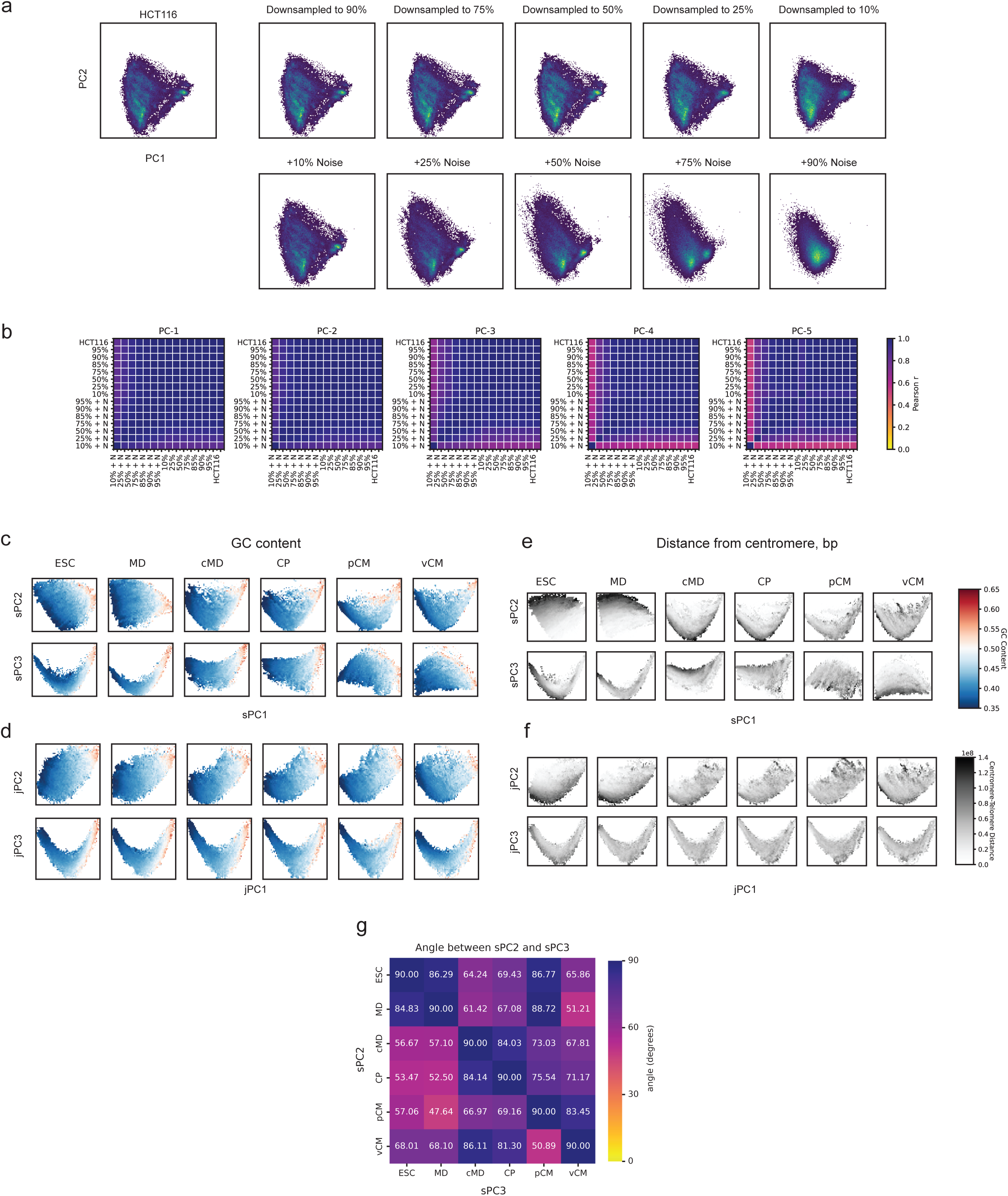
Sensitivity of jPCA to sequencing depth and noise, and comparative analysis of sPCA and jPCA. (A) Scatter plots of PC1 vs PC2 scores for joint PCA of a 50-kb resolution HCT116 contact map before and after progressive downsampling of read pairs (top row) and downsampling with addition of simulated random ligation pairs up to the original read depth (bottom row). (B) Heatmaps of Pearson correlation coefficients between PC score vectors of the perturbed HCT116 datasets for PCs 1-5. (C) PC score scatter plots for samples representing six successive *in vitro* cardiomyocyte differentiation stages, derived by separate PCA (sPCs), and colored by GC content. Top row: sPC1 vs sPC2 for each stage. Bottom row: sPC1 vs sPC3 for each stage. (D) Same as (C) but for PC scores derived by joint PCA (jPCs). Top: jPC1 vs jPC2 for each stage. Bottom row: jPC1 vs jPC3 for each stage. (E) Same as (C) but colored by genomic distance from the centromere. (F) Same as (D) but colored by genomic distance from the centromere. (G) Heatmap of matrix of acute angles in degrees between sPC2 and sPC3 basis vectors across differentiation stages.

**S.Figure 2.**
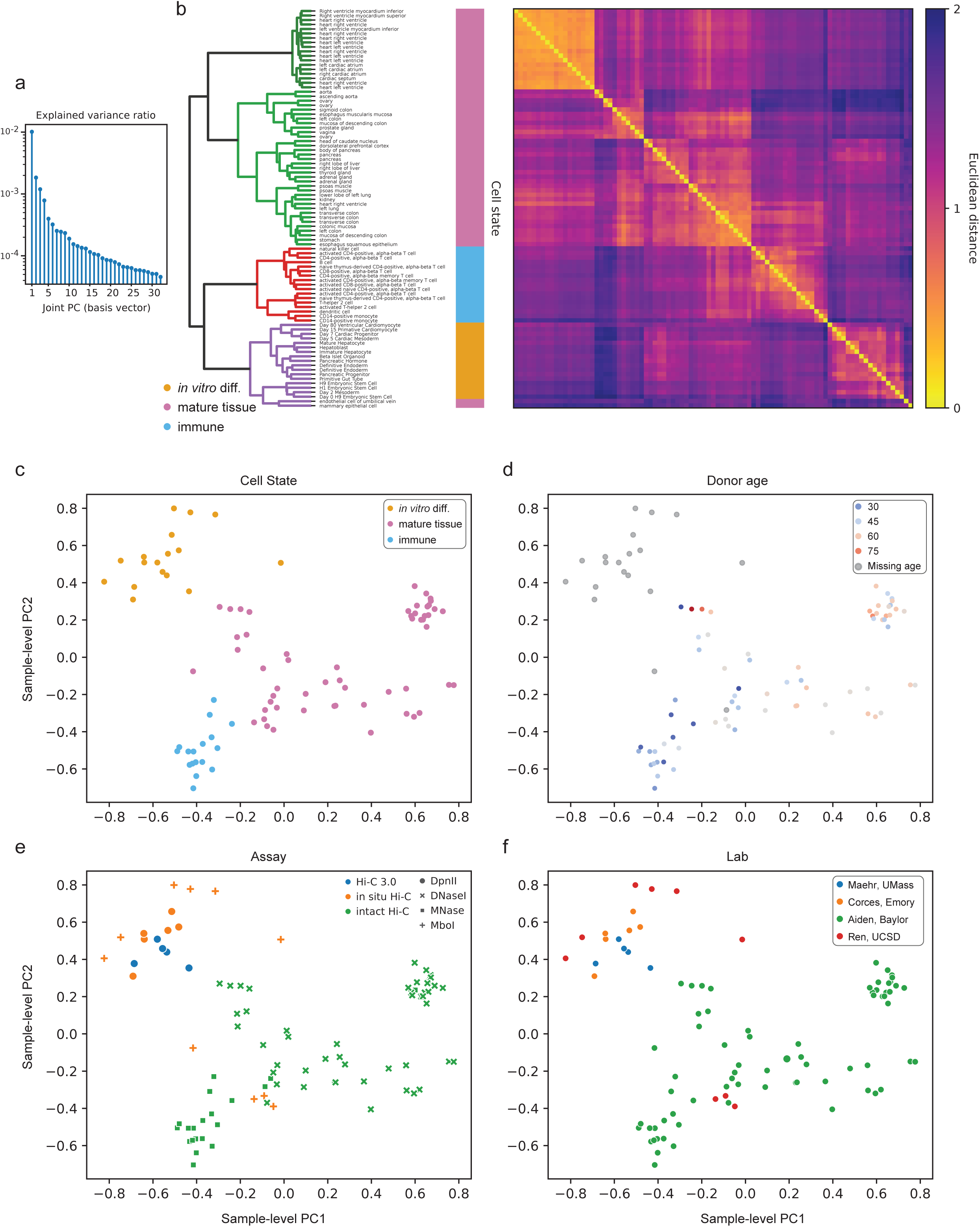
Sample-level hierarchical clustering and similarity of genome-wide contact frequency profiles as measured by joint PCA. (A) Explained variance ratios of atlas PCs. (B) Ward clustering dendrogram and euclidean distance heatmap between sample-level embeddings of the first 12 PC score vectors. (C-F) Sample-level PCA plot as in Figure 2B colored by sample covariates including: (C) assigned cell state category, (D) donor age, (E) Hi-C assay protocol, and (F) production lab.

**S.Figure 3.**
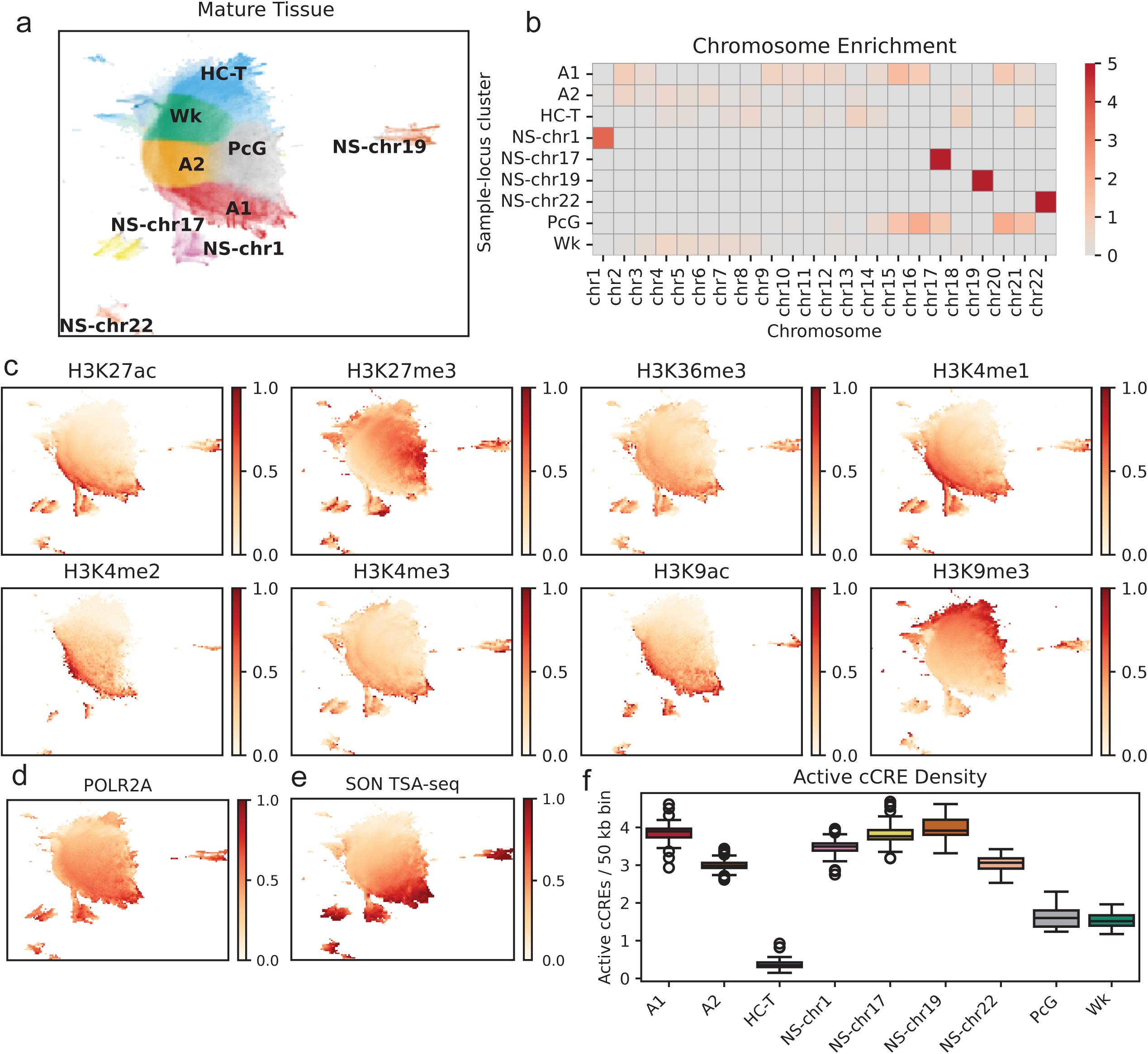
Four sample-locus clusters correspond to nuclear speckle-associated regions on specific chromosomes. (A) Visualization of a UMAP embedding all genomic bins from 55 mature tissue samples colored by sample-locus cluster label. NS-chr1, NS-chr17, NS-chr19, and NS-chr22 correspond to chromosome-specific nuclear speckle “islands”. B) Chromosome-level enrichment for sample-locus clusters in mature tissue samples (observed / expected) shows that speckle-associated island clusters are enriched for a specific chromosome. C) Mean ChIP-seq signal quantile from matched biosamples on ENCODE overlaid over UMAP embeddings. D) POLR2A ChIP-seq and E) SON TSA-seq mean signal quantiles overlaid against all bin embeddings show enrichment in A1 and speckle island clusters. F) Box and whisker plot of active cCREs per genomic bin across matching mature tissue biosamples for each sample-locus cluster.

**S.Figure 4.**
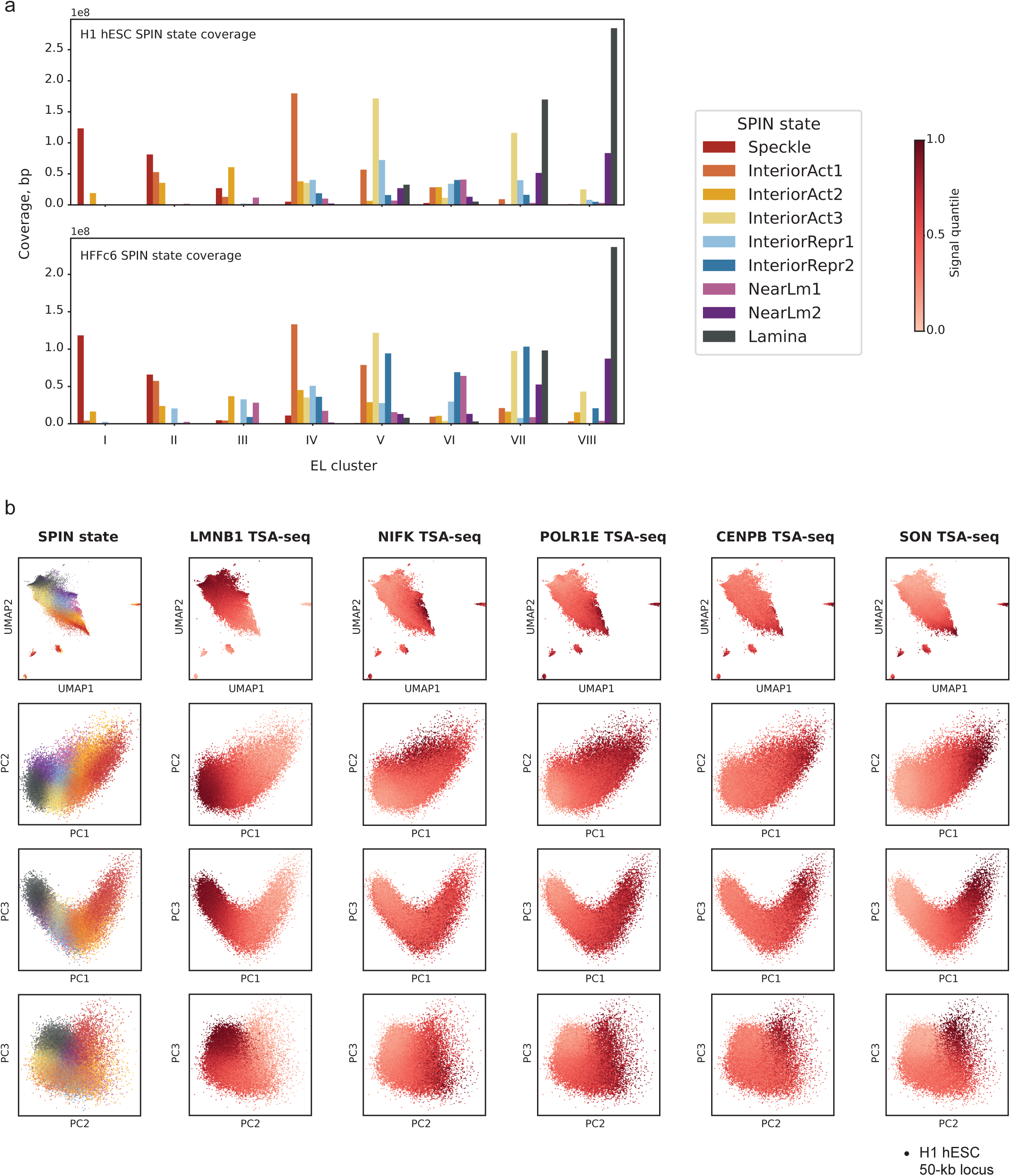
The spatial proximity to nuclear landmarks as measured by TSA-seq separates along multiple PCs. (A) Genomic coverage of SPIN state annotations from (top) H1 hESC and (bottom) HFFc6 cells overlapping ensemble-wide locus clusters. (B) Scatter plot visualizations of interaction profile UMAP embeddings and PCA projections of H1 hESC 50-kb loci colored by H1 SPIN state and by signal quantile of TSA-seq targeting marker proteins associated with the nuclear lamina (LMNB1), nucleolus (NIFK and POLR1E), centromere (CENPB), and nuclear speckles (SON) in H1 cells.

**S.Figure 5.**
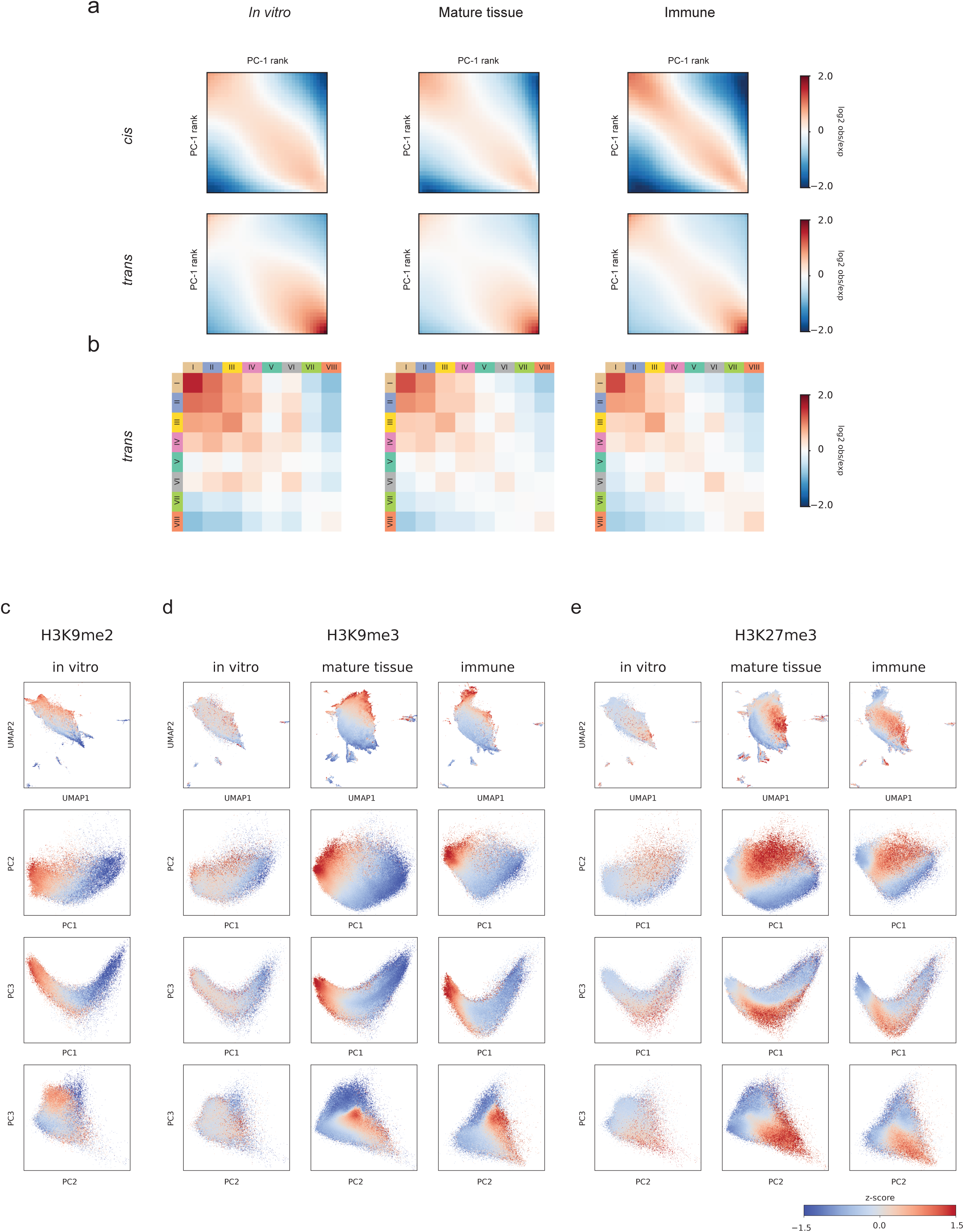
Saddle plots and locus embedding visualizations of repressive histone marks within each cell state category. (A) Continuous saddle plots for PC-1 rank in *cis* and *trans* for each cell state category. (B) Discrete saddle plots of EL clusters. (C-E) UMAP and PCA scatter plots of sample-locus long-range interaction embeddings colored by (C) H3K27me3, (D) H3K9me3, and (E) H3K9me2 signal z-score from matched biosamples for each cell state category.

**S.Figure 6.**
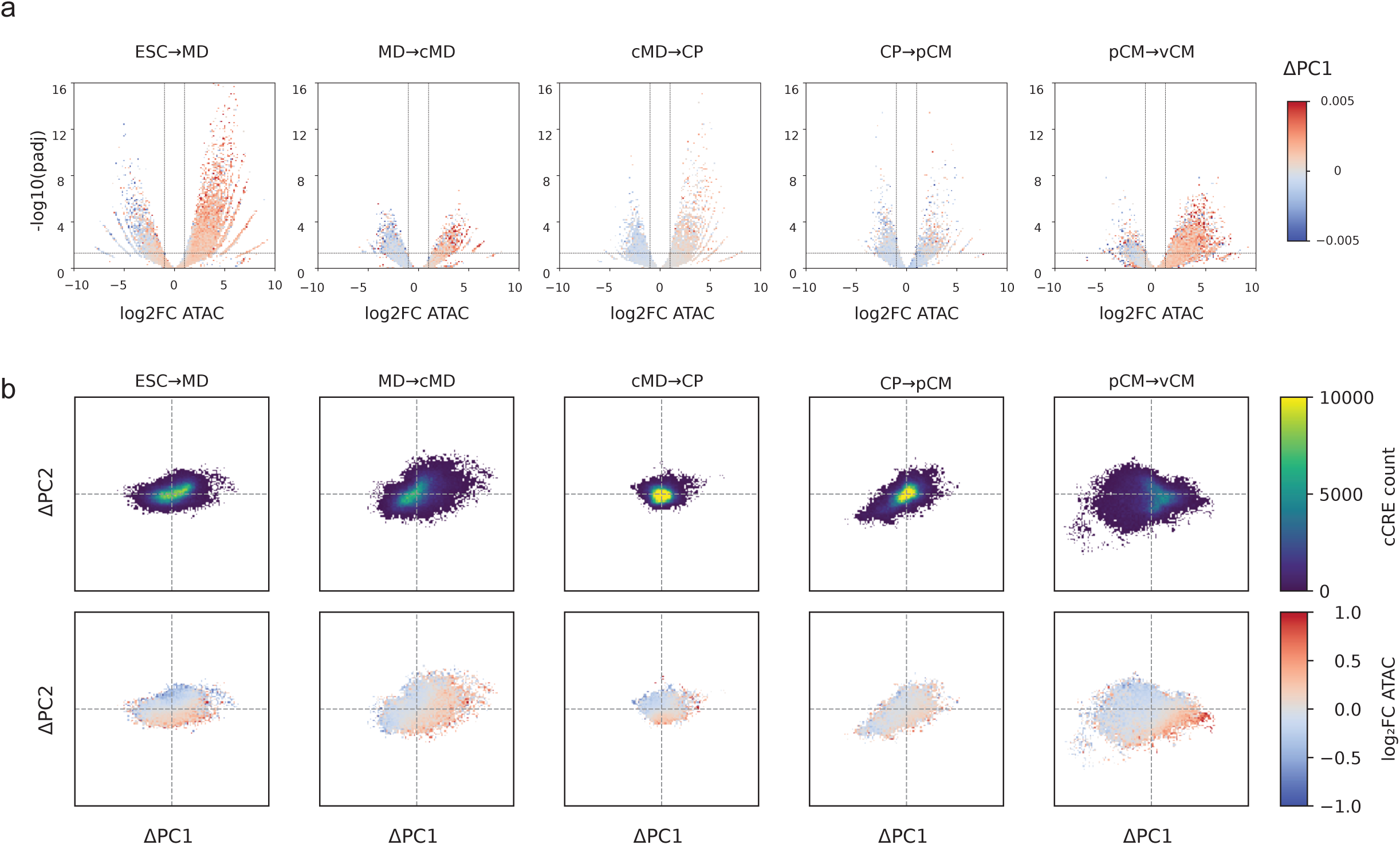
Relationship between accessibility of candidate *cis*-regulatory elements and changes in PC1 and PC2 during *in vitro* differentiation. (A) Volcano plots of differential chromatin accessibility at ENCODE cCREs between consecutive stages of cardiomyocyte differentiation, colored by change in PC1 score. (B) Scatter plots of change in PC1 score vs change in PC2 score at ENCODE cCREs, colored by point density (top) and log_2_ fold change ATAC-seq signal (bottom).

